# Genome-wide identification of functional tRNA-derived fragments in Senescence-accelerated mouse prone 8 brain

**DOI:** 10.1101/348300

**Authors:** Shuai Zhang, Hejian Li, Ling Zheng, Hong Li, Chengqiang Feng, Wensheng Zhang

## Abstract

tRNA-derived fragments (tRFs) have been linked previously to the development of various diseases, such as cancer and viral infection. However, tRFs seem also related to brain aging and related diseases, especially Alzheimer and Parkinson disease. RNA sequencing, a state-of-the-art technology, has allowed for investigation of tRFs in this field. In this study, we investigated the changes of tRFs in the brains of a senescence-accelerated mouse model, senescence-accelerated mouse prone 8 (SAMP8), that show age-dependent deficits in learning and memory; and a control model, senescence-accelerated mouse resistant 1 (SAMR1), with normal aging, both at 7 months of age. A total of 570 tRF transcripts were discovered. Among these transcripts, 8, including 3 upregulated and 5 downregulated transcripts, were differentially expressed in the SAMP8 mice. Then, we obtained 110 potential target genes in a miRNA-like pattern. GO survey implicated these target genes in the function of various aspects, e.g. postsynaptic density (GO: 0014069). Furthermore, we assessed in detail those tRFs whose miRNA-like pattern was most likely to affect the progression of either Alzheimer and Parkinson disease, such as AS-tDR-011775 acting on *Mobp* and *Park2*. In fact, we found the tRFs to be involved in the regulation of gene expression by means other than the miRNA-like pattern. Therefore, these 8 dysregulated tRFs may hold consequences far into the future and can be attractive biomarkers and valid targets. In brief, our study is the first to provide a comprehensive analysis on tRFs in SAMP8 mouse brain, and this breakthrough identified promising new targets for preventing the age-related changes of brain and the therapeutic intervention of Alzheimer’s and Parkinson’s.

## Introduction

Changes occur in all parts of a person’s body as the grows older, including the brain. The brain naturally shrinks in volume and there is increased size of the brain sulci with age (Raz and Rodrigue 2006). These changes have significant impacts on learning and other complex mental activities. It means that brain aging and neurodegeneration appear to go hand in hand, especially Alzheimer’s disease (AD) and Parkinson’s disease (PD) (Peters 2006; Duncan 2011). Brain aging-related diseases bring great agony and inconvenience to the patient’s life. Some countries, such as China, Japan, and Italy, have become aging societies that have considered the problem on aiding the aging brain and prevention of related diseases a public concern. Although many theories, including Aβ deposition and tau phosphorylation in AD (Swerdlow 2007; Alavi *et al*. 2015), and alpha-synuclein aggregation in PD (Xu and Pu 2016), have attempted to explain their origin, few treatment options are available to prevent the situation. Currently, gene expression regulation and brain aging and related diseases have drawn increasing attention among researchers.

In recent years, it has become apparent that the non-coding RNAs (ncRNAs) are of crucial functional importance for gene expression regulation. For example, as post transcriptional regulatory molecules that interact with specific mRNAs. These ncRNAs include miRNAs, piRNAs, snoRNAs, lncRNAs, circRNAs and so on. ncRNAs and their associated orchestrated networks are implicated in gene expression and mediating complex mechanisms of brain aging and related diseases. Some examples: circulatory miR-34a is an accessible biomarker for age-dependent changes in brain (Li *et al*. 2011); In a research review, investigators showed that piRNAs may be involved in age-dependent histone control of complex networks of memory-related genes (Earls *et al*. 2014). As high-throughput sequencing technology advances, a novel class of small noncoding RNAs (ncRNAs) derived from tRNAs was detected and called tRNA-derived fragments (tRFs), with lengths ranging from 14-36 nucleotides (nt) (Kumar *et al*. 2016; Keam and Hutvagner 2015). They can be classified into tRF-5, tRF-3, tRF-1, i-tRF, and tiRNA (tiRNA-3 and tiRNA-5) (Lee *et al*. 2009; Pliatsika *et al*. 2017). Many studies have shown that tRFs have specific biological roles and are implicated in mediating complex pathological processes of diverse illnesses, such as cancer and viral infectious disease (Zhou *et al*. 2017; Yeung *et al*. 2009; Guzman *et al*. 2015). A well-studied research revealed that tRFs are associated with mammalian brain aging (Karaiskos and Grigoriev 2016). tRFs have the potential effect in neurodegenerative processes (Hanada *et al*. 2013; Ivanov *et al*. 2014). However, our understanding of brain aging associated tRFs is limited only on preliminary explorations. The role of tRFs in AD and PD (two main types of brain aging-related diseases) is poorly known, at least, to date.

The senescence-accelerated mouse prone 8 (SAMP8) has about half the normal lifespan of a rodent and displays the early onset of senility accompanied by significant age-related deteriorations in memory and learning ability (Akiguchi *et al*. 2017). It is thus a potential useful mouse model to study brain aging and related disorders, particularly the application of the SAMP8 mice to AD and PD research (Porquet *et al*. 2013; Liu *et al*. 2008). The senescence-accelerated mouse resistant 1 (SAMR1) strain undergoes a normal aging process and is often used as the reference group (Zhang *et al*. 2008). In our previous studies, we analyzed the changes of lncRNA, circRNA, miRNA, and DNA methylation in the brain of SAMP8, and the related mechanisms for regulating gene expression (Zhang *et al*. 2016; Zhang *et al*. 2017; Zhang *et al*. 2017). Are there any changes of tRFs in SAMP8 mouse brain? How these changes affect the brain aging process and pathologic pattern of AD and PD? Given this, we characterized the expression of tRFs in the brain of SAMP8 and SAMR1 mice at 7 months of age through deep RNA sequencing in this study. Our research is the first to provide systematic insight into the profiling of the tRFs transcriptome in the brain aging model SAMP8 mouse. These tRFs may be potential therapeutic targets and diagnostic markers in brain aging associated illnesses, mainly AD and PD.

## Results

### Memory impairments at the 7-month stage in the SAMP8 mice

Morris water maze (MWM) test was performed to evaluate the learning and memory deficits in 7-month-old SAMP8 mice. The result for the hidden platform test was shown in **Figure 1A**; the SAMP8 mice entailed a longer time to find the platform than that of the SAMR1 mice (*p* < 0.05). The spatial probe test was performed next. **Figure 1B** clearly illustrated that the SAMR1 mice searched for the destination location purposefully, whereas the SAMP8 mice swam aimlessly in the pool. The number of crossings and the time percentage in the target quadrant were significantly lower in the SAMP8 group than in the SAMR1 group (*p* < 0.05, **Figures 1C and 1D**). When it comes to swimming speed, there had no difference between the two groups. (*p* > 0.05, **Figure 1E**); this result suggests the lack of motor and visual dysfunction in the SAMP8 mice. Evidently, the 7-month-old SAMP8 mice presented impaired memory and poor learning skills, which were consistent with clinical neurophysiology of aging brain and related neurodegeneration clinical symptoms.

**Figure 1.**
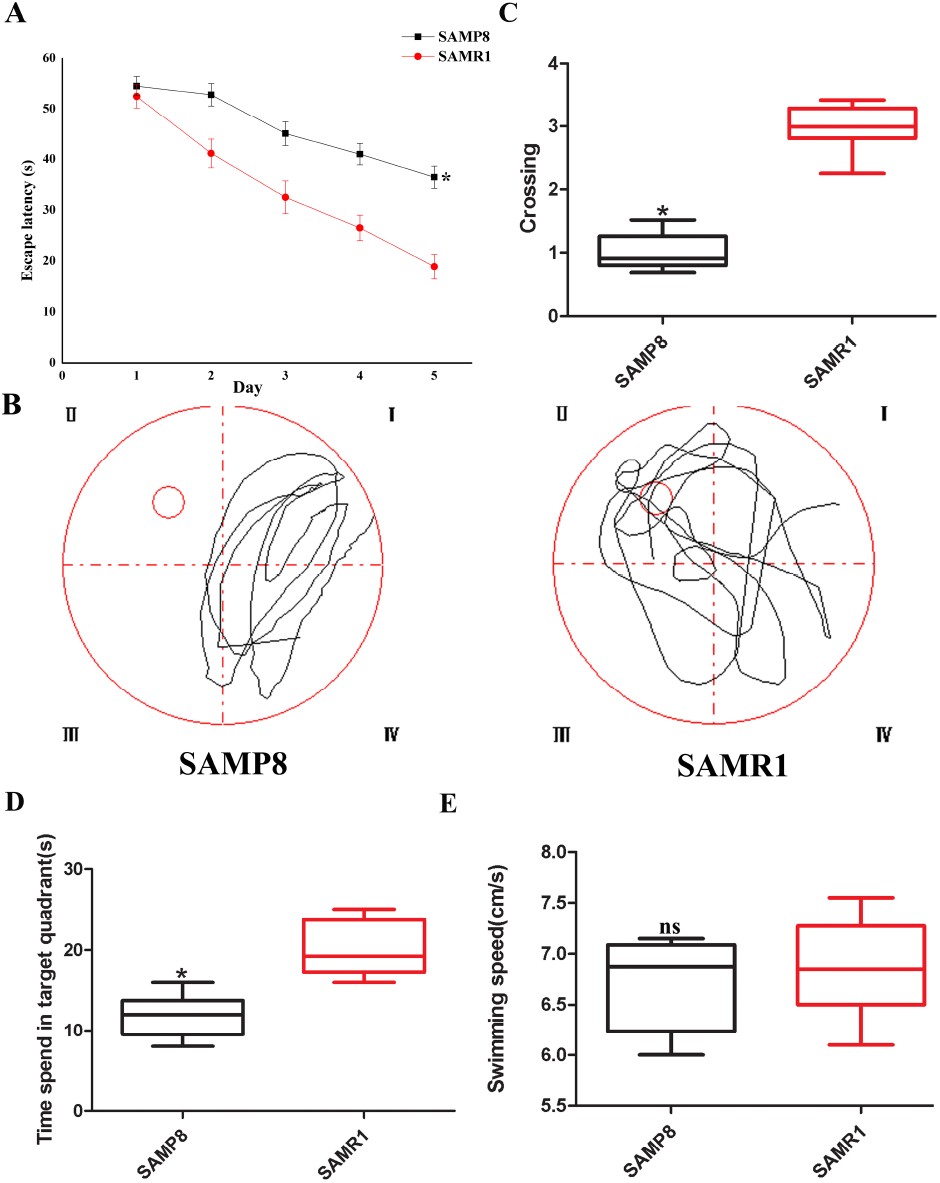
Memory impairments in SAMP8 mice. MWM test was used to evaluate the learning and memory deficits in 7-month-old SAMP8 and SAMR1 mice (n = 8/group). (A) Mean escape latency in the hidden platform experiment. (B) Swimming paths in the spatial probe experiment. (C) Number of crossings in the spatial probe experiment. (D) Time spent in the target quadrant in the spatial probe experiment. (E) The swimming speeds of the mice were similar between the two groups. \**p* < 0.05, ns means non-significant.

### tRFs sequencing roundup

A total of 69,772,438 raw reads (34,909,558 for the SAMP8 mice and 34,862,880 for SAMR1 mice) were generated. After the 5′- and 3′-adaptors were trimmed, low-quality reads were removed, and ≤16 bp reads were filtered, a total of 68,118,335 clean reads (33,886,463 for SAMP8 mice and 34,231,872 for SAMR1 mice) were found in the two groups. Most clean reads were 22, 21, 23, and 45 nt in length for both groups (**Supplementary Figures S1A and S1B**). Then, the high-quality clean data were mapped to the mouse mature-tRNA and pre-tRNA sequences from GtRNAdb by NovoAlign software (v2.07.11). In accordance with the comparison results, 570 tRFs were detected. These tRFs were used for subsequent analysis.

### Dysregulation of several tRFs in the SAMP8 mouse brain

We used transcripts per million (TPM) to estimate the expression level of the tRF transcripts. The expression level of each subtype showed a similar proportion between the two groups. The percentages were approximately 45% tRF-5, 26% tiRNA (2% tiRNA-3 and 24% tiRNA-5), 19% i-tRF, 5% tRF-3, and 5% tRF-1 (**Figures 2A and 2B**). As a result, 13 significantly differentially expressed tRFs were identified (*p* < 0.01 and fold changes ≥ 2). To validate the changes of detected by RNA-seq, all 13 tRFs were selected and their expression was further examined by qPCR. As shown in Figure 3, the results revealed that all 13 transcripts were detected in SAMP8 and SAMR1 brain, of which 8 were showed differential expression (*p* < 0.01, **Supplementary Table S1**). 5 was not shown differential expression which was inconsistent with RNA-seq data. This could be because the biological differences between samples.

**Figure 2.**
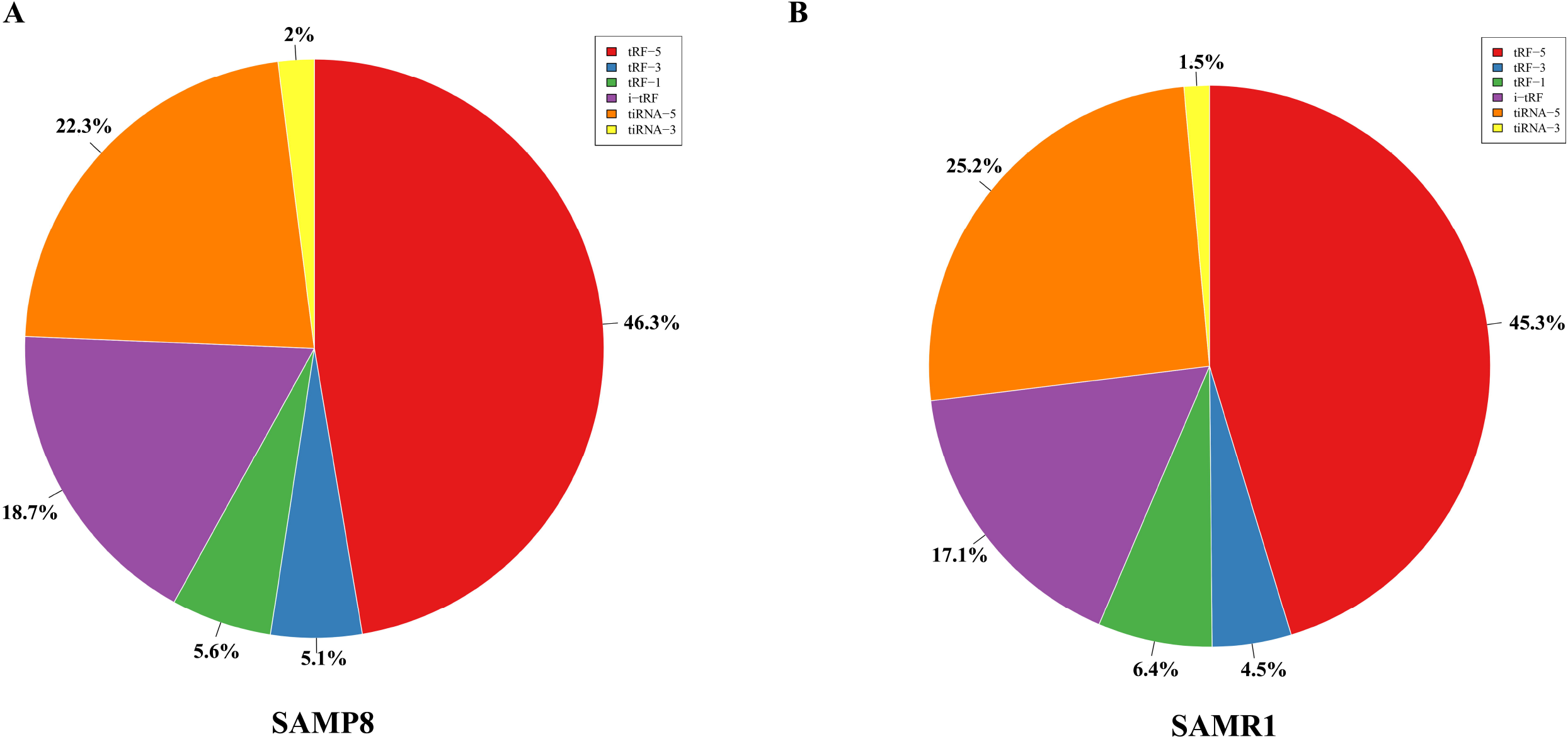
Proportions of tRF-5, tiRNA, i-tRF, tRF-3, and tRF-1 in the two groups. (A) Proportions in the SAMP8 mice. (B) Proportions in the SAMR1 mice.

**Figure 3.**
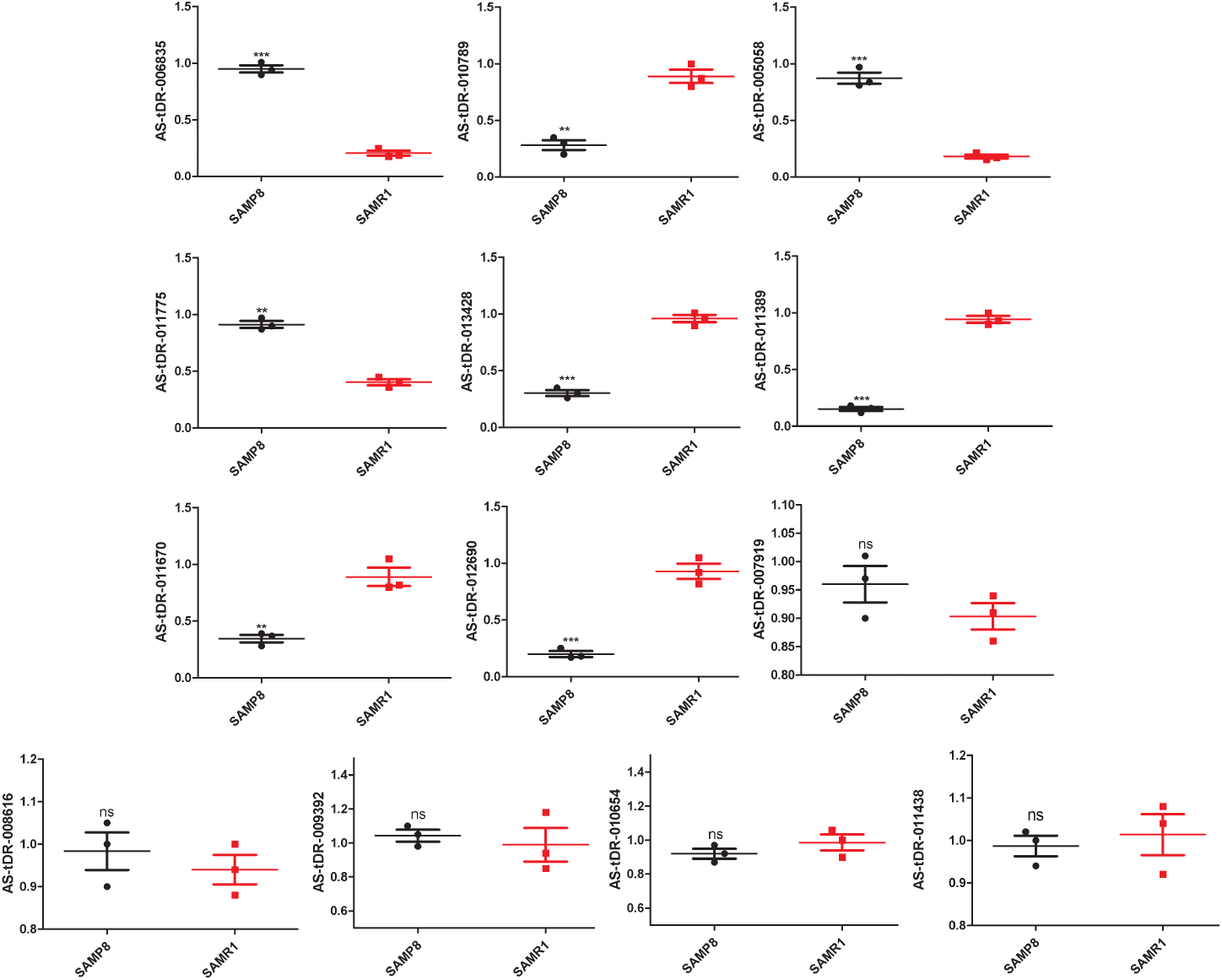
Validation of dysregulated tRFs by qPCR between SAMP8 and SAMR1 mice. The U6 gene was used as a housekeeping internal control. The relative expression of each tRF was represented as mean ± SEM [n = 3, three mice per group (biological replicates), three times per mouse (technical replicates)]. *p < 0.05, **p < 0.01, ***p < 0.001, ns means non-significant.

Then, cluster analysis of the 8 differently expressed genes tRFs was conducted with a heat map (**Figure 4**). In the SAMP8 group, three samples were clustered together. The same situation occurred in the SAMR1 group.

**Figure 4.**
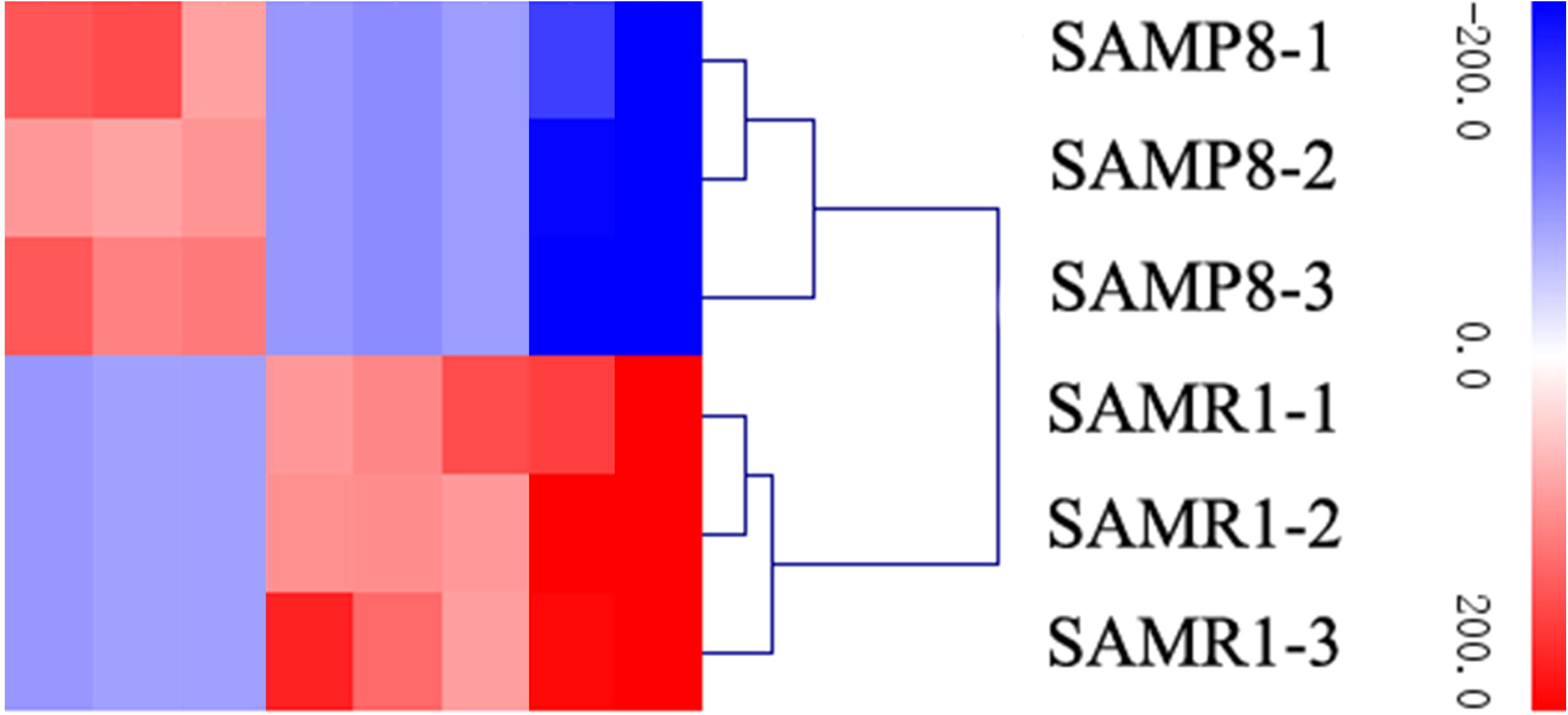
Cluster analysis of differentially expressed tRFs through a heat map. Red indicates an increased expression, and blue indicates a decreased expression.

### Target gene

We proposed that tRFs played extremely important roles in controlling gene transcription and translation by various means. Among them, a highly important pattern is the miRNA-like behavior (Karaiskos *et al*. 2015; Kumar *et al*. 2014). On the basis of this concept, we pioneered the identification of tRF-mRNA pairs in the SAMP8 brain through our mRNA-seq and tRFs-seq data. We chose to use 482 mRNAs and 8 tRFs that were differentially expressed in this study. The results are presented in **Supplementary Table S2**; 110 potential target genes were identified.

### GO enrichment

GO survey was performed on the above-mentioned target genes. As a result, 168 GO terms were significantly enriched (adjusted *p* value < 0.01, **Supplementary Table S3**). Importantly, several brain function associated terms were detected, including postsynaptic membrane (GO: 0045211), postsynaptic density (GO: 0014069), and neurogenesis (GO: 0022008). Overall, these protein-coding genes may be regulated by tRFs using a miRNA-like pattern involved in the SAMP8 mice brain.

### Relevance of the research

To obtain an improved understanding of the relationship between tRFs and brain aging and related diseases, especially AD and PD, we set three restrictions. For the first factor, the tRFs and their target genes must be expressed differently between the SAMP8 and SAMR1 mice. For the second factor, the trend of expression of tRF and its target gene should be the opposite in the brain. These trends include two cases: one case corresponds to the tRF (upregulated in the SAMP8 mice)-mRNA (downregulated in the SAMP8 mice), and the other case refers to the tRF (downregulated in the SAMP8 mice)-mRNA (upregulated in the SAMP8 mice). For the last requirement, the selected pairs (tRF-mRNA) should possess a close relationship with pathologic process of AD and PD. If a pair meets the above criteria, it can be selected. Take for example: Camk2n1, an endogenous CaMKII inhibitor protein, showed a direct effect on synaptic CaMKII-NMDAR binding and played an important role in LTP regulation (Gouet *et al*. 2012). We found that the expression of *Camk2n1* in the SAMP8 mice brain was significantly higher than that in the SAMR1 mice. AS-tDR-011389 presented the low expression level in the SAMP8 mouse brain and targeted *Camk2n1*. More specific details were listed in **Table 1**. We predicted that these tRFs most likely participate in the occurrence and development of AD and PD.

**Table 1.**
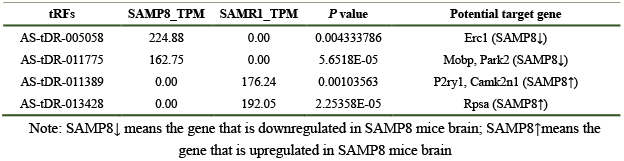
tRFs are most possibly involved in the pathogenesis of two main types of brain aging-related diseases (AD and PD) through a miRNA-like pattern

| tRFs | SAMP8_TPM | SAMR1_TPM | P value | Potential target gene |
| --- | --- | --- | --- | --- |
| AS-tDR-005058 | 224.88 | 0.00 | 0.004333786 | Erc1 (SAMP8↓) |
| AS-tDR-011775 | 162.75 | 0.00 | 5.6518E-05 | Mobp, Park2 (SAMP8↓) |
| AS-tDR-011389 | 0.00 | 176.24 | 0.00103563 | P2ry1, Camk2n1 (SAMP8↑) |
| AS-tDR-013428 | 0.00 | 192.05 | 2.25358E-05 | Rpsa (SAMP8↑) |

Note: SAMP8↓ means the gene that is downregulated in SAMP8 mice brain; SAMP8↑ means the gene that is upregulated in SAMP8 mice brain

Herein, we reiterate that besides the miRNA-like pattern, tRFs can also exert regulatory functions in other manners. For example, tRFs displaced the eIF4G translation initiation factor from mRNAs and participated in post-transcriptional regulation (Ivanov *et al*. 2011). The tRFs were capable of competing for the mRNA binding sites of YBX1 to suppress cancer progression (Goodarzi *et al*. 2015). Therefore, the functions of these tRFs (8 significant differentially expressed tRFs) may hold important meanings for the anti-brain aging and restraining neurodegenerative diseases.

## Discussion

In year 2009, Cole *et al*. first identified tRFs from cultured HeLa cells (Cole *et al*. 2009). Then, tRFs were detected in other kinds of human cells or tissues (Zhou *et al*. 2017; Yeung *et al*. 2009; Wang *et al*. 2013). tRFs were also evaluated in plants or other animals, such as barley (Hackenberg *et al*. 2013) and cattle (Casas *et al*. 2015). Given their widespread presence, the tRFs are expected to play key roles in many physiological and pathological processes. With aging, tRFs undergo dynamic changes in the mammalian brain (Karaiskos and Grigoriev 2016). Getting older underlies cognitive decline and dementia, and is the most dangerous element for the failure of brain function. The study of aging brain can shed light on the basic neurological mechanisms and their connections with age-related neurodegenerative conditions, such as AD and PD. AD is the most common neurodegenerative disorder worldwide and the leading cause of dementia among older adults (Jahn 2013; Fratiglion and Qiu 2009). PD is the second most common neurodegenerative disorder, after AD (Spillantini *et al*. 1998). They are two major types of brain aging-related neurodegenerative disorders. No valid therapeutic options are available to prevent or reverse them. Finding early diagnostic biomarkers and novel therapeutic targets is thus imperative. Recent studies have shifted focus to dysregulated gene alterations and regulations in brain aging and related diseases, particularly AD and PD. We have conducted considerable work previously on this aspect, including lncRNA, miRNA, circRNA, and DNA methylation (Zhang *et al*. 2016; Zhang *et al*. 2017; Zhang *et al*. 2017). The anomaly of tRFs may exert a great influence on brain aging and related diseases. As a special and relative new ncRNA, tRFs may be correlated with brain aging and related diseases development and advance by regulating the expression of specific genes. To date, the research on this aspect is quite limited. Additional research and attention should be paid to it. In the present study, the primary goal is to identify tRFs which are involved in brain aging and related diseases as much as possible. Unfortunately, the human tissue samples are very difficult to collect for several reasons. For this reason, we selected an idealized animal model to break past the limit. Research indicates that the SAMP8 mice exhibit age-related brain degeneration and AD and PD like pathologies (Kang *et al*. 2014; Liu *et al*. 2008). Our MWM test confirmed that the SAMP8 mice exhibited memory impairments and learning deficits at the 7-month-old; these signs were the core clinical features observed in older people. Moreover, high-throughput sequencing (RNA-seq) was applied in our study and allowed the identification of potential tRFs in unprecedented detail. On the basis of the results, the candidate tRFs can be explored as novel sensitive factors for improving brain aging and AD and PD control in the future.

First, we discovered 570 tRF transcripts in SAMP8 and SAMR1 mouse brains at 7 months of age. On the basis of the standard classification (Lee *et al*. 2009), these tRFs were classified into various subtypes, including 69 tiRNA-5, 14 tiRNA-3, 94 tRF-3, 94 tRF-5, 67 tRF-1, and 232 i-tRF. Karaiskos *et al*. described the age-driven modulation of tRFs in Drosophila but focused specifically on the tRF species containing CCA at the 3’-end (Karaiskos *et al*. 2015). In the current study plenty of 5′-derived tRFs (tRF-5 and tiRNA-5) and i-tRFs have also been discovered in aging brain. tiRNA-5, tiRNA-3, tRF-3, tRF-5, and i-tRF series which are generated from mature tRNAs constituted the majority in our samples, whereas the tRFs (tRF-1) generated from the primary tRNAs were in the minority. This phenomenon is consistent with previous reports (Casas *et al*. 2015; Olvedy *et al*. 2016; Wei *et al*. 2012).

Second, we calculated the expression of each tRF transcript through TPM. Although i-tRF was the subtype with the most tRF transcripts, the expression levels of the i-tRF transcripts were not the highest. The tRF-5 subtype achieved the highest expression levels. On the opposite end, the expression levels of the tiRNA-3 was the lowest. The 5’-derived tRFs (tRF-5 and tiRNA-5) were the most abundant class of tRFs between the two groups, followed by i-tRF, with the 3′-derived tRFs (tiRNA-3, tRF-3 and tRF-1) least.

Third, the tRFs with fold changes ≥ 2 and *p* < 0.01 were selected as differential expression in this sequencing. A total of 13 tRF transcripts were preliminary found. Then, qPCR was used to detect the accuracy and reliability of the initial results. Ultimately, 8 dysregulated tRFs were detected, including 3 upregulated and 5 downregulated tRFs in SAMP8 mice relative to those in SAMR1 mice. This finding demonstrated that our pipeline was relatively high-quality strict in identifying putative tRFs and laid a solid foundation for further exploration and experimentation. Briefly, AS-tDR-011775, AS-tDR-006835 and AS-tDR-005058 expressed significantly higher in SAMP8 mice brain than that in SAMR1 mice brain. By contrast, AS-tDR-013428, AS-tDR-011389, AS-tDR-012690, AS-tDR-010789 and AS-tDR-011670 expression were significantly higher in SAMR1 mice brain. Through cluster analysis, we hypothesized the relationships among samples. In our study, the result of cluster analysis showed a distinguishable tRF expression profiling between the two groups. We were aware that tRFs are tied closely to various diseases (Fu *et al*. 2015; Anderson and Ivanov 2014). It is predicted that these dysregulated tRFs probably play functional roles in brain aging and related diseases.

Fourth, tRFs are proven to play key roles in the process of gene expression and regulation in several manners. A fundamental path is to behave similar to miRNAs (Karaiskos *et al*. 2015; Kumar *et al*. 2014). Given previous experience, the interactions of the protein-coding genes and tRFs were determined through miRanda and TargetScan. We then identified 110 potential genes. Through GO survey, these target genes were found to be involved in brain function from different aspects, such as neurogenesis (GO: 0022008), postsynaptic density (GO: 0014069), and postsynaptic membrane (GO: 0045211). Furthermore, several special filter conditions were set to select the most likely tRF-mRNA pairs involved in the regulation of brain aging and the most underlying illnesses (AD and PD) progression under a miRNA-like pattern. We mentioned the AS-tDR-011389-Camk2n1 pair in previous text (Gouet *et al*. 2012). *Rpsa*, a gene found to facilitate the production and internalization of neurotoxic Aβ peptide (Jovanovic *et al*. 2015), was targeted by AS-tDR-013428. AS-tDR-011775 acted on *Mobp. Mobp* may play a role in controlling axonal diameter and keeping axons round (Sadahiro *et al*. 2000). Park2 has greater relevance to Parkinson’s disease (Nuytemans *et al*. 2010). *Park2* was targeted by AS-tDR-011775. P2Y1 receptor protein is encoded by the *P2ry1* gene. A study confirmed that P2Y1 receptor mediated the astroglial network dysfunction by purinergic signaling in AD (Delekate *et al*. 2014). *P2ry1* was regulated by AS-tDR-011389. AS-tDR-005058 acted on *Erc1*. Study showed that neurotransmitter release can be regulated through the Rab6ip2/ERC1/ CAST2/ELKS and presynaptic active zone RIM proteins interaction (Wang *et al*. 2002). Interestingly, one tRF can dominate an increased number of genes. AS-tDR-011389 is a good example and controlled *P2ry1* and *Camk2n1*. This observation suggests that the miRNA-like mechanism of tRFs in the regulation of gene expression is complex in brain aging. Moreover, the participation of tRFs in gene expression regulation by other means must be emphasized. tRFs are capable of displacing the eIF4G translation initiation factor from mRNAs (Ivanov *et al*. 2011). tRFs suppress breast cancer progression via YBX1 displacement (Goodarzi *et al*. 2015). These tRFs (8 dysregulated tRFs) may hold important implications in the anti-brain aging and control and prevention of neurodegenerative diseases. Therefore, our ongoing effort will focus on the functions of these high-potential tRFs at the molecular level. This goal is expected to be a massive challenge for many years to come.

To summarize, our study investigated tRF profiles in the brain of SAMP8 and SAMR1 mice at 7 months’ age and offered a lead for the further investigation of the biological role and marker potential of tRFs in brain aging and related diseases, especially AD and PD. The study of tRFs in this respect has merely commenced but will certainly open a new frontier in the development of new therapeutic targets for anti-brain aging and control and reduce the currence of relevant diseases in the future.

## Materials and Methods

### Preparation of animals

In this study task, we purchased SAMP8 mice (n = 15, 3 months of age, male, pathogen and virus free, RRID: MGI: 2160863) and SAMR1 mice (n = 15, 3 months of age, male, pathogen and virus free, RRID: MGI: 2160867) from Beijing WTLH Biotechnology Co., Ltd. (Licence No.SCXK Jing 2011-0012). The mice were maintained in separate cages with standard conditions and allowed to obtain food and water freely until 7 months old. No animals died and were excluded. All were used in the following experiments. Eight animals of each group were randomly selected for the Morris water maze (MWM) test, numbered 1 to 8. The remaining mice were given isoflurane anesthesia (isoflurane is a nonflammable liquid administered by vaporizing, is a general inhalation anesthetic drug. Briefly: first, place the animal in the induction chamber; second, adjust the flowmeter to 0.8 to 1.5 L/min; third, adjust the isoflurane vaporizer to 3%-5%), euthanized by cervical dislocation, and dissected to obtain their cerebral cortices. The tissues were immediately preserved in liquid nitrogen at −196 °C for tRF sequencing and other experiments.

All experimentation on the mice complied with the “Guide for the care and use of laboratory animals” (National Research Council 2010) and were permitted by the Institutional Animal Care and Use Committee of Beijing Normal University (BNU NO. 2018)

### Behavioral studies

The spatial learning and memory of SAMP8 mice at the 7-month-old stage were evaluated through the MWM as previously described (Zhang *et al*. 2017). Briefly, the mice were familiarized with the MWM environment on the day before the program. In the hidden platform experiment (days 1-5), we set a platform in a suppositive quadrant. The mice were trained twice a day for 5 days. The mice were then allowed to swim for 90 s during each training. The escape latency was recorded through a special software once they touched the platform. However, if a mouse failed to reach the platform within the stipulated time, we helped find the platform, and the escape latency was regarded as 90 s. In the spatial probe experiment (day 6), we withdrew the platform and allowed the mice to swim freely for 1 min. The time spent in the target quadrant, the number of platform crossing, and the swimming trajectory of each mouse within 1 min were recorded. All experiments were performed simultaneously every day, and the investigator was unaware of the mouse genotypes throughout the trial.

### Library preparation

Six cDNA libraries were constructed, i.e., three for the SAMP8 mice and three for the SAMR1 mice. Agarose gel electrophoresis was adopted to examine the integrality of total RNA samples and were quantified on the NanoDrop ND-2000 instrument (Thermo Scientific™, USA, #ND-2000). Total RNA samples were first pretreated as follows to remove some RNA modifications that interfere with small RNA-seq library construction: 3′-aminoacyl deacylation to 3′-OH for 3′ adaptor ligation, 3′-cP removal to 3′-OH for 3′ adaptor ligation, 5′-OH phosphorylation to 5′-P for 5′-adaptor ligation, m1A and m3C demethylation for efficient reverse transcription. These steps were conducted to prepare a gene library: 1) 3′-adapter ligation, 2) 5′-adapter ligation, 3) cDNA synthesis, 4) PCR amplification, and 5) size selection of ~135-160 bp PCR-amplified fragments (corresponding to ~15-40 nt small RNAs). The Agilent bioanalyzer 2100 system (Agilent, USA, #G2939BA) was used to assess library quality. Finally, the libraries were pooled in equal amounts depending on the quantification results.

### Sequencing

The libraries were denatured and diluted to a loading volume of 1.3 ml and loading concentration of 1.8 pM with 0.1 M NaOH. The diluted libraries were then loaded onto a reagent cartridge and forwarded to a sequencing run on the Illumina NextSeq 500 system (RRID: SCR_014983) by using a NextSeq 500/550 V2.5 kit (Illumina, USA, #FC-404) in accordance with manufacturer’s instructions. Sequencing was carried out in 50 cycles.

### Quality control and mapping summary

Raw data files in FASTQ format were generated through Illumina NextSeq 500. The sequencing quality was shown by quality score. This quality score was then represented by Q, which is the −10×log10 transformed probability of the base calling being incorrect. Q30 means the incorrect probability is 0.001. If the number is larger than 30, the incorrect probability is less than 0.001, i.e., > 99.9% correct. Generally, when most of the quality scores are above 30, the sequence is of high quality.

After Illumina quality control, the sequencing reads were 5′,3′-adaptor trimmed, filtered for ≥ 16 nt by the Cutadapt software (Martin 2011), and aligned to mature-tRNA and pre-tRNA sequences from GtRNAdb (Chan and Lowe 2016) by using the NovoAlign software (v2.07.11) (Wang *et al*. 2016).

### Expression analysis

tRF expression levels were measured and normalized as read counts per million of total aligned tRF reads (TPM). tRFs with fold changes ≥ 2 and *p* < 0.01 were selected as the significantly differentially expressed tRFs between SAMP8 and SAMR1.

### Quantitative real-time PCR

The results of tRFs-seq were validated through qPCR. qPCR was performed with the ViiA7 Real-time PCR System, rtStar™ tRF&tiRNA Pretreatment Kit (Arraystar, USA, #AS-FS-005), rtStar™ First-Strand cDNA Synthesis Kit (Arraystar, USA, # AS-FS-003), and 2×PCR master mix (Arraystar, USA, #AS-MR-005). The specific quantitative primers are listed in **Table 2**. The 10 μL reaction volume contained 0.5 μL of each primer, 2 μL of H_2_O, 2 μL of cDNA, and 5 μL of 2× Master Mix. The conditions were 95 °C for 10 min followed by 40 cycles (95 °C for 10 s and 60 °C for 60 s). Each experiment was performed in triplicate.

**Table 2.**
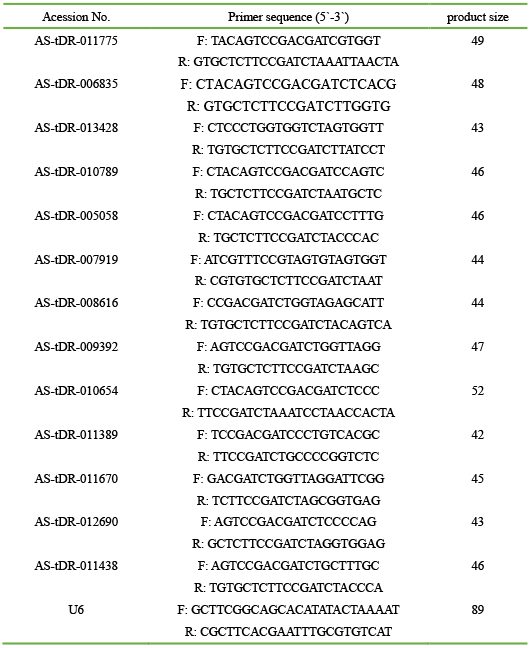
Primers used in qPCR analysis

### Target prediction

Research has shown that a highly important function of tRFs is to behave like miRNAs and repress the expression of endogenous targets (Karaiskos *et al*. 2015; Kumar *et al*. 2014). In other words, the tRFs pair with the 3′UTRs of the mRNAs to direct the latter’s post-transcriptional repression. Given this observation, we used miRanda and TargetScan to systematically predict the tRF-mRNA interaction. In this research, the expression levels of the tRFs and mRNAs showed significant difference between the SAMP8 and SAMR1 mice and were hence analyzed. Our previous study (Zhang *et al*. 2016) revealed that 482 mRNA transcripts were differentially expressed between SAMP8 and SAMR1 mouse brains at 7 months of age (adjusted *p* value < 0.05, **Supplementary Table S4**).

### Gene Ontology (GO) survey

GO enrichment analysis was applied to the target genes of tRFs. GOseq R package was then used to perform GO analysis (Young *et al*. 2010). GO terms with adjusted *p* value < 0.01 were recognized as significant enrichment.

### Statistical analysis

The test results were analyzed by SPSS 20.0 and Graph pad prism 5 software. Box plot was conducted to show distribution of data into quartiles. The ends of the box are the upper and lower quartiles. The median is marked by a vertical line inside the box. The whiskers are the two lines outside the box that extend to the highest and lowest observations. Dot plot was conducted to show qPCR data. *p* < 0.05 represented significant difference. The difference of the escape latency data in the MWM test was compared with two-way ANOVA. Student’s t test was applied to compare the qPCR results, and the remaining data of the MWM test.

## Acknowledgments

This work was supported by the National Nature Science Foundation of China(81771152), the National Key Research and Development Plan of China (2017YFC1702500).We also thank Mr Hui Ma and his colleagues at Kangcheng Biotech for their assistance.

## Author contributions

SZ WZ conceived and designed the experiments. SZ HL LZ HL performed the experiments. SZ HL CF analyzed the data. WZ contributed to the acquisition of reagents/materials/analysis tools. SZ wrote the paper.

## Data availability

The tRFs-seq raw data have been deposited in the NCBI Sequence Read Archive (SRA). The accession number is PRJNA533967.

## Preprint

There is a preprint at https://www.biorxiv.org/content/early/2018/06/15/348300 (Zhang *et al*. 2018).

## Abbreviations

AD: Alzheimer’s disease
GO: Gene Ontology
MWM: Morris water maze
ncRNAs: Non-coding RNAs
PD: Parkinson’s disease
RRID: Research Resource Identifier
SAMP8: Senescence-accelerated mouse prone 8
SAMR1: Senescence-accelerated mouse resistant 1
TPM: Transcripts per million
tRFs: tRNA-derived fragments

## Supplementary material

**Supplementary Figure S1. Sequence length distribution of clean reads in the two groups**. (A) Sequence length distribution in the SAMP8 mice. (B) Sequence length distribution in the SAMR1 mice.

**Supplementary Table S1**. Significantly and differentially expressed tRF transcripts between SAMP8 and SAMR1 mice.

**Supplementary Table S2**. The potential targets of tRFs.

**Supplementary Table S3**. GO enrichment analysis of the tRF-targeting genes.

**Supplementary Table S4**. Significantly and differentially expressed mRNA transcripts between SAMP8 and SAMR1 mice.

