## Supplementary figures and images for "Genome-wide identification of functional tRNA-derived fragments in Senescence-accelerated mouse prone 8 brain"

### Supplemental Figure 1

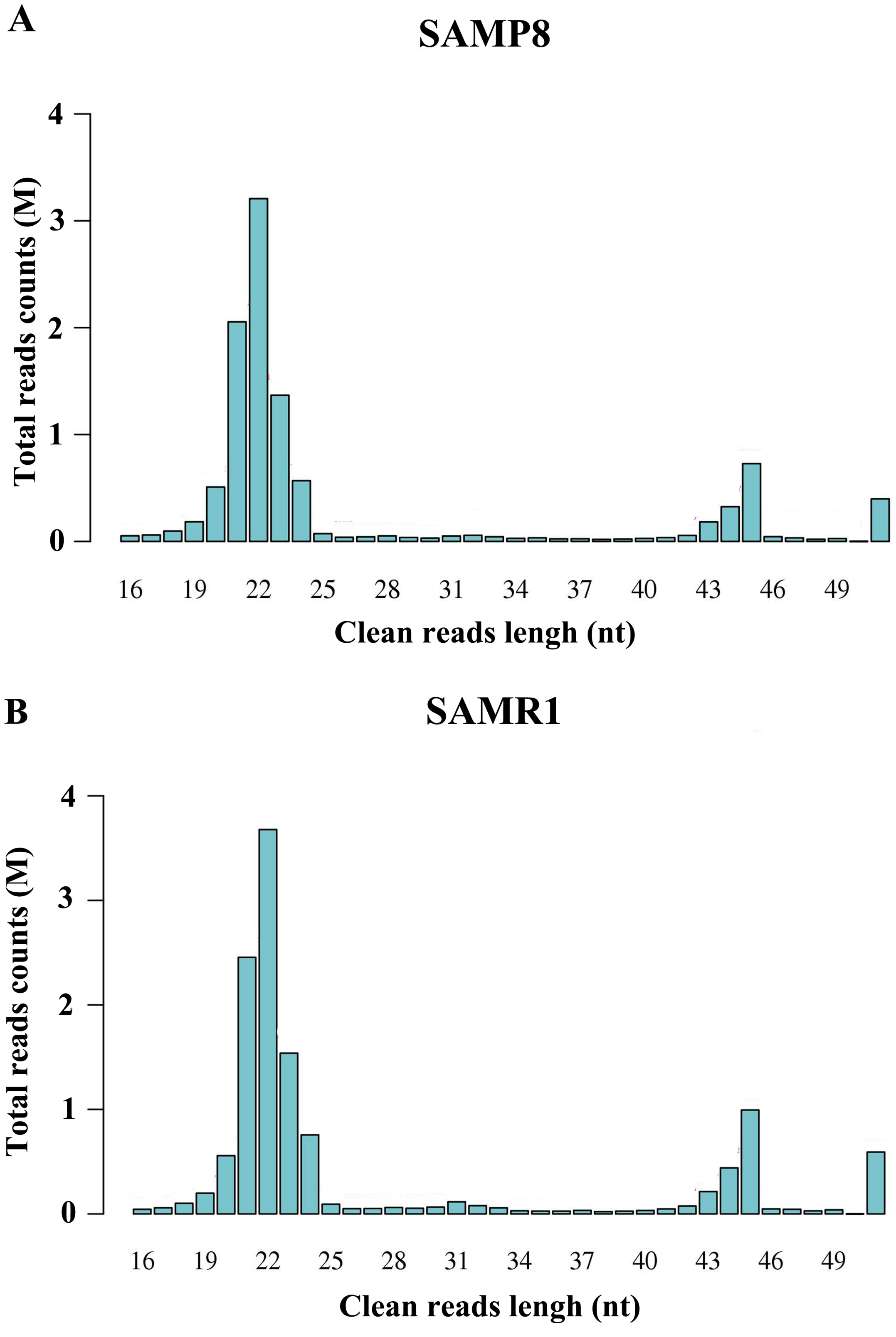
