## Supplemental Table 1 for "Genome-wide identification of functional tRNA-derived fragments in Senescence-accelerated mouse prone 8 brain"

**Table S2 Significantly and differentially expressed tRF transcripts between SAMP8 and SAMR1 mice**

| tRFs_ID | source tRNA | Left | Right | tRF_Length | tRF_Sequence | SP8_TPM | SR1_TPM | fold_change | *p* value |
|---|---|---|---|---|---|---|---|---|---|
| AS-tDR-011775 | pre-VAl-TAC-1-1 | 114 | 132 | 19 | GTGGTGTGCTAGTTAATTT | 162.75 | 0.00 | 163 | 5.6518E-05 |
| AS-tDR-006835 | Trp-CCA-1-1 | 59 | 75 | 17 | TCACGTCGGGGTCACCA | 179.04 | 0.00 | 179 | 0.000780738 |
| AS-tDR-005058 | Ser-GCT-3-1 | 48 | 63 | 16 | CTTTGCACGCGTGGGT | 204.88 | 0.00 | 205 | 0.004333786 |
| AS-tDR-013428 | Glu-CTC-2-1 | 1 | 26 | 26 | TCCCTGGTGGTCTAGTGGTTAGGATA | 0.00 | 192.05 | -192 | 2.25358E-05 |
| AS-tDR-010789 | Lys-TTT-1-1 | 13 | 28 | 16 | CAGTCGGTAGAGCATT | 200.89 | 572.08 | -2.84 | 0.000124317 |
| AS-tDR-011389 | Asp-GTC-2-1 | 31 | 53 | 23 | CCTGTCACGCGGGAGACCGGGGC | 0.00 | 176.24 | -176 | 0.00103563 |
| AS-tDR-012690 | Ala-AGC-3-1 | 58 | 75 | 18 | TCCCCAGCATCTCCACCT | 0.00 | 196.14 | -196 | 0.002818642 |
| AS-tDR-011670 | Glu-CTC-1-1 | 16 | 42 | 27 | TGGTTAGGATTCGGCGCTCTCACCGCT | 665.27 | 1731.64 | -2.6 | 0.004289344 |
