## Supplemental Table 2 for "Genome-wide identification of functional tRNA-derived fragments in Senescence-accelerated mouse prone 8 brain"

**Table S3 The potential targets of tRFs**

| tRFs | Target transcript_id | Target gene_name |
|---|---|---|
| AS-tDR-011389 | ENSMUST00000029632 | Lrat |
| AS-tDR-012690 AS-tDR-013428 | ENSMUST00000090180 | Sema3g |
| AS-tDR-012690 AS-tDR-011389 | ENSMUST00000110690 | Oxct1 |
| AS-tDR-012690 | ENSMUST00000187137 | Mag |
| AS-tDR-013428 | ENSMUST00000086545 | Cdkl4 |
| AS-tDR-012690 | ENSMUST00000102518 | Ece1 |
| AS-tDR-013428 AS-tDR-011670 | ENSMUST00000178401 | Zfp870 |
| AS-tDR-011775 | ENSMUST00000077741 | Slc9a6 |
| AS-tDR-012690 | ENSMUST00000153941 | Slc38a6 |
| AS-tDR-011775 | ENSMUST00000191124 | Park2 |
| AS-tDR-011775 AS-tDR-013428 | ENSMUST00000033342 | Eif3f |
| AS-tDR-012690 | ENSMUST00000060792 | Ptrf |
| AS-tDR-012690 | ENSMUST00000192145 | Lrba |
| AS-tDR-010789 AS-tDR-011389 | ENSMUST00000006669 | Pdk1 |
| AS-tDR-012690 | ENSMUST00000164960 | Rasgef1a |
| AS-tDR-013428 | ENSMUST00000062387 | Kcnj9 |
| AS-tDR-012690 | ENSMUST00000067230 | Sox4 |
| AS-tDR-011389 | ENSMUST00000068261 | Atp6v1g2 |
| AS-tDR-012690 | ENSMUST00000022416 | Anxa11 |
| AS-tDR-012690 | ENSMUST00000109815 | Camk2b |
| AS-tDR-011389 | ENSMUST00000047309 | Nat14 |
| AS-tDR-013428 | ENSMUST00000189707 | Ppp2r3d |
| AS-tDR-011775 | ENSMUST00000102852 | Ptges |
| AS-tDR-012690 | ENSMUST00000021203 | Timm22 |
| AS-tDR-011389 | ENSMUST00000101800 | Msrb1 |
| AS-tDR-011389 | ENSMUST00000005620 | Dnajb1 |
| AS-tDR-011389 | ENSMUST00000045356 | Rpl37 |
| AS-tDR-011389 | ENSMUST00000050918 | Camk2n1 |
| AS-tDR-011775 | ENSMUST00000174193 | Mobp |
| AS-tDR-011670 AS-tDR-006835 | ENSMUST00000082432 | Dio2 |
| AS-tDR-011775 | ENSMUST00000062181 | Zfp146 |
| AS-tDR-010789 AS-tDR-012690 | ENSMUST00000006618 | Arhgef19 |
| AS-tDR-013428 | ENSMUST00000145910 | Strn |
| AS-tDR-013428 | ENSMUST00000172753 | Hspa1b |
| AS-tDR-011775 | ENSMUST00000140119 | Col9a2 |
| AS-tDR-011389 | ENSMUST00000057486 | Ankrd46 |
| AS-tDR-011670 | ENSMUST00000034121 | Man2b1 |
| AS-tDR-005058 | ENSMUST00000055389 | Xxylt1 |
| AS-tDR-011670 AS-tDR-012690 | ENSMUST00000034713 | Ldlr |
| AS-tDR-011775 | ENSMUST00000059650 | Npsr1 |
| AS-tDR-010789 | ENSMUST00000030029 | Invs |
| AS-tDR-012690 | ENSMUST00000133544 | Tor1b |
| AS-tDR-010789 | ENSMUST00000027343 | Ogfrl1 |
| AS-tDR-011670 | ENSMUST00000154650 | Bcas1 |
| AS-tDR-013428 | ENSMUST00000035105 | Rpsa |
| AS-tDR-011389 | ENSMUST00000051867 | Lsm6 |
| AS-tDR-005058 | ENSMUST00000118012 | Gm14295 |
| AS-tDR-013428 | ENSMUST00000102484 | Ddi2 |
| AS-tDR-011389 | ENSMUST00000092227 | Scyl2 |
| AS-tDR-012690 | ENSMUST00000042844 | Nbl1 |
| AS-tDR-011389 | ENSMUST00000029331 | P2ry1 |
| AS-tDR-011389 | ENSMUST00000159879 | Trove2 |
| AS-tDR-011670 | ENSMUST00000163582 | Ptp4a3 |
| AS-tDR-012690 | ENSMUST00000112640 | Steap3 |
| AS-tDR-012690 | ENSMUST00000080322 | Oas1a |
| AS-tDR-012690 AS-tDR-013428 | ENSMUST00000045224 | Acer2 |

| | | |
|---|---|---|
| AS-tDR-012690 | ENSMUST00000020957 | Adi1 |
| AS-tDR-011670 AS-tDR-012690 AS-tDR-013428 | ENSMUST00000015449 | Sash1 |
| AS-tDR-013428 | ENSMUST00000099294 | Ablim1 |
| AS-tDR-011670 | ENSMUST00000045537 | Chrm4 |
| AS-tDR-011775 | ENSMUST00000048718 | Mmaa |
| AS-tDR-013428 | ENSMUST00000065103 | Mrpl35 |
| AS-tDR-012690 | ENSMUST00000167323 | Apold1 |
| AS-tDR-011775 | ENSMUST00000061852 | Dclre1c |
| AS-tDR-013428 | ENSMUST00000063976 | Opa3 |
| AS-tDR-012690 | ENSMUST00000046285 | C1qa |
| AS-tDR-012690 | ENSMUST00000053020 | Neurl1b |
| AS-tDR-011775 AS-tDR-013428 | ENSMUST00000066708 | Dmp1 |
| AS-tDR-005058 | ENSMUST00000069180 | Zcchc24 |
| AS-tDR-011775 | ENSMUST00000039205 | Galm |
| AS-tDR-013428 | ENSMUST00000151224 | Fam163b |
| AS-tDR-012690 | ENSMUST00000127748 | Tril |
| AS-tDR-013428 | ENSMUST00000109626 | Cyld |
| AS-tDR-011670 AS-tDR-011775 | ENSMUST00000045376 | Adk |
| AS-tDR-011389 AS-tDR-011670 | ENSMUST00000055738 | Tsc22d3 |
| AS-tDR-012690 | ENSMUST00000024884 | Eif2ak2 |
| AS-tDR-012690 | ENSMUST00000090287 | Myh11 |
| AS-tDR-005058 | ENSMUST00000118009 | Naf1 |
| AS-tDR-010789 AS-tDR-013428 AS-tDR-01269 | ENSMUST00000171588 | Lifr |
| AS-tDR-005058 | ENSMUST00000113975 | Slc5a3 |
| AS-tDR-013428 | ENSMUST00000111194 | Cd44 |
| AS-tDR-005058 | ENSMUST00000045756 | S100a10 |
| AS-tDR-011775 | ENSMUST00000098822 | Zfp606 |
| AS-tDR-012690 AS-tDR-013428 | ENSMUST00000169927 | Adora1 |
| AS-tDR-012690 | ENSMUST00000060214 | Tmem125 |
| AS-tDR-010789 AS-tDR-012690 | ENSMUST00000106252 | Mycl |
| AS-tDR-012690 | ENSMUST00000205741 | Nelfb |
| AS-tDR-011389 | ENSMUST00000087328 | Hspa1a |
| AS-tDR-005058 | ENSMUST00000002176 | Celf2 |
| AS-tDR-005058 | ENSMUST00000105285 | Epyc |
| AS-tDR-012690 | ENSMUST00000106355 | Zfp691 |
| AS-tDR-011775 AS-tDR-013428 | ENSMUST00000075304 | Stx3 |
| AS-tDR-011389 | ENSMUST00000106635 | Kcnj16 |
| AS-tDR-005058 | ENSMUST00000053308 | Ranbp3l |
| AS-tDR-012690 | ENSMUST00000002379 | Cd320 |
| AS-tDR-005058 AS-tDR-012690 | ENSMUST00000040912 | Anln |
| AS-tDR-012690 | ENSMUST00000172815 | Gm19345 |
| AS-tDR-011775 | ENSMUST00000112186 | Mettl8 |
| AS-tDR-011389 | ENSMUST00000060716 | Samd3 |
| AS-tDR-011775 AS-tDR-012690 | ENSMUST00000038423 | Rtp4 |
| AS-tDR-013428 | ENSMUST00000020102 | Slc17a8 |
| AS-tDR-011389 | ENSMUST00000032198 | Usp18 |
| AS-tDR-005058 AS-tDR-013428 | ENSMUST00000032279 | Erc1 |
| AS-tDR-011389 | ENSMUST00000072271 | Etnppl |
| AS-tDR-011775 | ENSMUST00000085668 | Gm5113 |
| AS-tDR-013428 | ENSMUST00000057243 | Tmem252 |
| AS-tDR-012690 | ENSMUST00000081982 | Dzank1 |
| AS-tDR-011389 AS-tDR-012690 | ENSMUST00000068975 | Zfp180 |
| AS-tDR-011670 | ENSMUST00000098230 | Rhog |
| AS-tDR-005058 AS-tDR-011775 | ENSMUST00000077687 | Ccdc148 |
