## Supplemental Table 3 for "Genome-wide identification of functional tRNA-derived fragments in Senescence-accelerated mouse prone 8 brain"

**Table S4 GO enrichment analysis of the tRFs-targeting genes**

| GO accession | Description | adjusted *p* value |
|---|---|---|
| GO:0005488 | binding | 9.63368E-12 |
| GO:0009987 | cellular process | 7.02926E-11 |
| GO:0005737 | cytoplasm | 7.02926E-11 |
| GO:0005623 | cell | 1.6044E-10 |
| GO:0044464 | cell part | 1.6044E-10 |
| GO:0043226 | organelle | 1.66199E-10 |
| GO:0044699 | single-organism process | 5.11122E-10 |
| GO:0044424 | intracellular part | 1.21588E-09 |
| GO:0043229 | intracellular organelle | 1.33876E-09 |
| GO:0005622 | intracellular | 1.47705E-09 |
| GO:0043227 | membrane-bounded organelle | 2.5191E-09 |
| GO:0016020 | membrane | 2.90816E-09 |
| GO:0065007 | biological regulation | 7.54182E-09 |
| GO:0005515 | protein binding | 1.06246E-08 |
| GO:0050789 | regulation of biological process | 1.12978E-08 |
| GO:0044444 | cytoplasmic part | 2.25253E-08 |
| GO:0043231 | intracellular membrane-bounded organelle | 3.36657E-08 |
| GO:0044763 | single-organism cellular process | 4.72509E-08 |
| GO:0060076 | excitatory synapse | 1.18944E-07 |
| GO:0050896 | response to stimulus | 2.74024E-07 |
| GO:1902578 | single-organism localization | 7.80717E-07 |
| GO:0099572 | postsynaptic specialization | 7.80717E-07 |
| GO:0014069 | postsynaptic density | 7.80717E-07 |
| GO:0044456 | synapse part | 1.113E-06 |
| GO:0044425 | membrane part | 2.36004E-06 |
| GO:0008152 | metabolic process | 2.42575E-06 |
| GO:0044765 | single-organism transport | 3.12808E-06 |
| GO:0051179 | localization | 3.26027E-06 |
| GO:0071704 | organic substance metabolic process | 4.304E-06 |
| GO:0044238 | primary metabolic process | 7.31219E-06 |
| GO:0071840 | cellular component organization or biogenesis | 7.74386E-06 |
| GO:0032501 | multicellular organismal process | 8.5263E-06 |
| GO:0016021 | integral component of membrane | 1.51359E-05 |
| GO:0044237 | cellular metabolic process | 1.65959E-05 |
| GO:0045202 | synapse | 1.75704E-05 |
| GO:0044707 | single-multicellular organism process | 2.10428E-05 |
| GO:0031224 | intrinsic component of membrane | 2.7244E-05 |
| GO:0016043 | cellular component organization | 3.28013E-05 |
| GO:0051234 | establishment of localization | 5.82518E-05 |
| GO:0050794 | regulation of cellular process | 6.58827E-05 |
| GO:0071944 | cell periphery | 6.78208E-05 |
| GO:0006810 | transport | 9.08927E-05 |
| GO:0003824 | catalytic activity | 9.22741E-05 |
| GO:0098794 | postsynapse | 0.000112185 |
| GO:0048518 | positive regulation of biological process | 0.000120916 |
| GO:0005886 | plasma membrane | 0.000121196 |
| GO:0048584 | positive regulation of response to stimulus | 0.000122445 |
| GO:0071702 | organic substance transport | 0.000147776 |
| GO:0042221 | response to chemical | 0.000147776 |
| GO:0044260 | cellular macromolecule metabolic process | 0.000147776 |
| GO:0044710 | single-organism metabolic process | 0.000152126 |
| GO:0044459 | plasma membrane part | 0.00016792 |

| | | |
|---|---|---|
| GO:0033036 | macromolecule localization | 0.000167937 |
| GO:0043170 | macromolecule metabolic process | 0.000172734 |
| GO:0048731 | system development | 0.000191289 |
| GO:0005773 | vacuole | 0.000191289 |
| GO:0044700 | single organism signaling | 0.000191289 |
| GO:0023052 | signaling | 0.000194183 |
| GO:0048583 | regulation of response to stimulus | 0.00020298 |
| GO:0007154 | cell communication | 0.000237139 |
| GO:0097458 | neuron part | 0.000264179 |
| GO:0012505 | endomembrane system | 0.000295956 |
| GO:0030030 | cell projection organization | 0.000367791 |
| GO:0051716 | cellular response to stimulus | 0.000423236 |
| GO:0022008 | neurogenesis | 0.000443306 |
| GO:0097060 | synaptic membrane | 0.000457414 |
| GO:0048856 | anatomical structure development | 0.000457414 |
| GO:0003008 | system process | 0.000461325 |
| GO:0007275 | multicellular organism development | 0.000548502 |
| GO:0046983 | protein dimerization activity | 0.000577189 |
| GO:0051239 | regulation of multicellular organismal process | 0.000677565 |
| GO:0007399 | nervous system development | 0.000689487 |
| GO:0031175 | neuron projection development | 0.000746378 |
| GO:0007267 | cell-cell signaling | 0.000776767 |
| GO:0010647 | positive regulation of cell communication | 0.000780327 |
| GO:0048699 | generation of neurons | 0.000789999 |
| GO:0007165 | signal transduction | 0.000796339 |
| GO:0023056 | positive regulation of signaling | 0.000796339 |
| GO:0044281 | small molecule metabolic process | 0.000940652 |
| GO:0030182 | neuron differentiation | 0.00102578 |
| GO:0045211 | postsynaptic membrane | 0.001089569 |
| GO:0044767 | single-organism developmental process | 0.00111014 |
| GO:0010817 | regulation of hormone levels | 0.001224305 |
| GO:0032502 | developmental process | 0.001310671 |
| GO:1902253 | egulation of intrinsic apoptotic signaling pathway by p53 class mediat | 0.001424227 |
| GO:0043412 | macromolecule modification | 0.001534588 |
| GO:0031988 | membrane-bounded vesicle | 0.001534588 |
| GO:0036211 | protein modification process | 0.001644732 |
| GO:0006464 | cellular protein modification process | 0.001644732 |
| GO:1901615 | organic hydroxy compound metabolic process | 0.001855141 |
| GO:0032879 | regulation of localization | 0.001872633 |
| GO:0031982 | vesicle | 0.001977693 |
| GO:0098793 | presynapse | 0.002029822 |
| GO:0072657 | protein localization to membrane | 0.002057844 |
| GO:0032990 | cell part morphogenesis | 0.002075714 |
| GO:0071705 | nitrogen compound transport | 0.002134696 |
| GO:1901363 | heterocyclic compound binding | 0.002166946 |
| GO:0048666 | neuron development | 0.002166946 |
| GO:0006807 | nitrogen compound metabolic process | 0.002291926 |
| GO:0008104 | protein localization | 0.002559966 |
| GO:0005771 | multivesicular body | 0.00270834 |
| GO:0097159 | organic cyclic compound binding | 0.002725873 |
| GO:0007155 | cell adhesion | 0.002727709 |
| GO:0044267 | cellular protein metabolic process | 0.002789965 |
| GO:0022610 | biological adhesion | 0.002843672 |
| GO:0098590 | plasma membrane region | 0.003066073 |

| | | |
|---|---|---|
| GO:0009967 | positive regulation of signal transduction | 0.003066073 |
| GO:0007166 | cell surface receptor signaling pathway | 0.003165071 |
| GO:0044802 | single-organism membrane organization | 0.003220703 |
| GO:0043065 | positive regulation of apoptotic process | 0.00339653 |
| GO:0051641 | cellular localization | 0.003428487 |
| GO:0043068 | positive regulation of programmed cell death | 0.003563562 |
| GO:0030544 | Hsp70 protein binding | 0.003862304 |
| GO:0048522 | positive regulation of cellular process | 0.004126204 |
| GO:0023061 | signal release | 0.00420703 |
| GO:0048869 | cellular developmental process | 0.004328606 |
| GO:0010646 | regulation of cell communication | 0.004328606 |
| GO:0065008 | regulation of biological quality | 0.004375719 |
| GO:1905114 | cell surface receptor signaling pathway involved in cell-cell signaling | 0.004648643 |
| GO:0023051 | regulation of signaling | 0.004648643 |
| GO:0031072 | heat shock protein binding | 0.004780788 |
| GO:0010942 | positive regulation of cell death | 0.005028518 |
| GO:0050877 | neurological system process | 0.005311221 |
| GO:0032844 | regulation of homeostatic process | 0.005331941 |
| GO:0001609 | G-protein coupled adenosine receptor activity | 0.005666327 |
| GO:0019538 | protein metabolic process | 0.005861788 |
| GO:0045184 | establishment of protein localization | 0.006062686 |
| GO:0046879 | hormone secretion | 0.006097226 |
| GO:0006950 | response to stress | 0.006115185 |
| GO:0061024 | membrane organization | 0.00615872 |
| GO:0002793 | positive regulation of peptide secretion | 0.006172485 |
| GO:0048468 | cell development | 0.006172485 |
| GO:0048858 | cell projection morphogenesis | 0.006202775 |
| GO:0051649 | establishment of localization in cell | 0.006203276 |
| GO:0072331 | signal transduction by p53 class mediator | 0.006203276 |
| GO:0070062 | extracellular exosome | 0.006203276 |
| GO:0009914 | hormone transport | 0.00628183 |
| GO:0009628 | response to abiotic stimulus | 0.00628183 |
| GO:1903561 | extracellular vesicle | 0.0064482 |
| GO:0042447 | hormone catabolic process | 0.006485653 |
| GO:0043230 | extracellular organelle | 0.006485653 |
| GO:0036477 | somatodendritic compartment | 0.006750473 |
| GO:0004872 | receptor activity | 0.006750473 |
| GO:0060089 | molecular transducer activity | 0.006750473 |
| GO:0099501 | exocytic vesicle membrane | 0.00677767 |
| GO:0030672 | synaptic vesicle membrane | 0.00677767 |
| GO:0070887 | cellular response to chemical stimulus | 0.007061643 |
| GO:0097367 | carbohydrate derivative binding | 0.007706876 |
| GO:0009966 | regulation of signal transduction | 0.007706876 |
| GO:0030154 | cell differentiation | 0.007842158 |
| GO:0006468 | protein phosphorylation | 0.007852659 |
| GO:1901796 | regulation of signal transduction by p53 class mediator | 0.007852659 |
| GO:0010467 | gene expression | 0.007930928 |
| GO:0045785 | positive regulation of cell adhesion | 0.007959156 |
| GO:0006812 | cation transport | 0.007970948 |
| GO:0042312 | regulation of vasodilation | 0.008122936 |
| GO:0044297 | cell body | 0.00819057 |
| GO:0009058 | biosynthetic process | 0.008274392 |
| GO:0006066 | alcohol metabolic process | 0.008376476 |
| GO:0051967 | negative regulation of synaptic transmission, glutamatergic | 0.008574097 |

| | | |
|---|---|---|
| GO:0051085 | chaperone mediated protein folding requiring cofactor | 0.008574097 |
| GO:0040011 | locomotion | 0.008574097 |
| GO:0043167 | ion binding | 0.008742456 |
| GO:0016055 | Wnt signaling pathway | 0.009434182 |
| GO:0198738 | cell-cell signaling by wnt | 0.009519801 |
| GO:0006915 | apoptotic process | 0.009742171 |
| GO:0001973 | adenosine receptor signaling pathway | 0.009904811 |
| GO:0090150 | establishment of protein localization to membrane | 0.009904811 |
