## Supplemental Table 4 for "Genome-wide identification of functional tRNA-derived fragments in Senescence-accelerated mouse prone 8 brain"

**Table S1 Significantly and differentially expressed mRNA transcripts between SAMP8 and SAMR1 mice**

| transcript_id | gene_id | gene_name | length | SAMP8_FPKM | SAMR1_FPKM | log2(foldchange) | pvalue | Adjusted p value |
|---|---|---|---|---|---|---|---|---|
| ENSMUST00000024823 | ENSMUSG00000024026 | Glo1 | 1935 | 50.0226 | 113.402 | -1.18079 | 5.00E-05 | 0.00896 |
| ENSMUST00000098519 | ENSMUSG00000036712 | Cyld | 7954 | 33.7971 | 19.1444 | 0.819982 | 5.00E-05 | 0.00896 |
| ENSMUST00000025511 | ENSMUSG00000024608 | Rps14 | 683 | 1029.44 | 710.914 | 0.534109 | 5.00E-05 | 0.00896 |
| ENSMUST00000082407 | ENSMUSG00000064356 | mt-Atp8 | 204 | 19184.9 | 12216.9 | 0.651087 | 5.00E-05 | 0.00896 |
| ENSMUST00000022816 | ENSMUSG00000022205 | Sub1 | 3372 | 102.171 | 70.0828 | 0.543855 | 5.00E-05 | 0.00896 |
| ENSMUST00000022954 | ENSMUSG00000022332 | Khdrbs3 | 1887 | 125.893 | 97.9376 | 0.362262 | 5.00E-05 | 0.00896 |
| ENSMUST00000029632 | ENSMUSG00000028003 | Lrat | 5351 | 0.878417 | 0.0857949 | 3.35594 | 5.00E-05 | 0.00896 |
| ENSMUST00000076922 | ENSMUSG00000030921 | Trim30a | 3771 | 0.645472 | 1.19475 | -0.88828 | 5.00E-05 | 0.00896 |
| ENSMUST00000072739 | ENSMUSG00000064317 | Gm10146 | 448 | 15.9499 | 6.18774 | 1.36606 | 5.00E-05 | 0.00896 |
| ENSMUST00000040402 | ENSMUSG00000036398 | Ppp1r11 | 1599 | 46.0024 | 30.988 | 0.569999 | 5.00E-05 | 0.00896 |
| ENSMUST00000189457 | ENSMUSG00000101972 | Hist1h3i | 411 | 35.444 | 17.8487 | 0.989718 | 5.00E-05 | 0.00896 |
| ENSMUST00000025178 | ENSMUSG00000024319 | Vps52 | 3332 | 13.7885 | 9.3241 | 0.56443 | 5.00E-05 | 0.00896 |
| ENSMUST00000177591 | ENSMUSG00000096768 | Erdr1 | 776 | 9.41966 | 4.23912 | 1.15191 | 5.00E-05 | 0.00896 |
| ENSMUST00000090776 | ENSMUSG00000071478 | Hist1h2ad | 463 | 7.03945 | 1.70679 | 2.04418 | 5.00E-05 | 0.00896 |
| ENSMUST00000110690 | ENSMUSG00000022186 | Oxct1 | 3527 | 178.117 | 85.0701 | 1.0661 | 5.00E-05 | 0.00896 |
| ENSMUST00000023779 | ENSMUSG00000023034 | Nr4a1 | 2474 | 17.55 | 25.4761 | -0.537675 | 5.00E-05 | 0.00896 |
| ENSMUST00000188775 | ENSMUSG00000101355 | Hist1h3h | 411 | 25.3941 | 12.1053 | 1.06886 | 5.00E-05 | 0.00896 |
| ENSMUST00000187994 | ENSMUSG00000075602 | Ly6a | 955 | 10.0793 | 0.570602 | 4.14276 | 5.00E-05 | 0.00896 |
| ENSMUST00000026911 | ENSMUSG00000025804 | Ccr1 | 2831 | 0.105164 | 0.74094 | -2.81672 | 5.00E-05 | 0.00896 |
| ENSMUST00000105046 | ENSMUSG00000078249 | Hmga1-rs1 | 1626 | 2.6375 | 6.28017 | -1.25163 | 5.00E-05 | 0.00896 |
| ENSMUST00000081342 | ENSMUSG00000094777 | Hist1h2ap | 482 | 8.19827 | 3.39358 | 1.27251 | 5.00E-05 | 0.00896 |
| ENSMUST00000011407 | ENSMUSG00000011263 | Exoc3l2 | 2201 | 0.139825 | 0.622708 | -2.15493 | 5.00E-05 | 0.00896 |
| ENSMUST00000102983 | ENSMUSG00000064288 | Hist1h4k | 312 | 270.927 | 173.858 | 0.639999 | 5.00E-05 | 0.00896 |
| ENSMUST00000086545 | ENSMUSG00000033966 | Cdkl4 | 2931 | 9.65715 | 5.08034 | 0.926672 | 5.00E-05 | 0.00896 |
| ENSMUST00000128727 | ENSMUSG00000056367 | Actr3b | 2031 | 36.5892 | 18.3164 | 0.998284 | 5.00E-05 | 0.00896 |
| ENSMUST00000122452 | ENSMUSG00000031698 | Mylk3 | 3036 | 1.0389 | 0.210241 | 2.30495 | 5.00E-05 | 0.00896 |
| ENSMUST00000064391 | ENSMUSG00000052560 | Cpne8 | 3366 | 4.41192 | 6.39133 | -0.534708 | 5.00E-05 | 0.00896 |
| ENSMUST00000116527 | ENSMUSG00000047844 | Bex4 | 837 | 19.359 | 9.10215 | 1.08872 | 5.00E-05 | 0.00896 |
| ENSMUST00000024572 | ENSMUSG00000023806 | Rsph3b | 1614 | 1.14776 | 0.455806 | 1.33233 | 5.00E-05 | 0.00896 |
| ENSMUST00000186219 | ENSMUSG00000072774 | Zfp951 | 2904 | 0.193768 | 0.772844 | -1.99585 | 5.00E-05 | 0.00896 |
| ENSMUST00000088940 | ENSMUSG00000038141 | Tmem181a | 4405 | 35.846 | 46.9413 | -0.389045 | 5.00E-05 | 0.00896 |
| ENSMUST00000099683 | ENSMUSG00000075014 | Gm10800 | 665 | 15.3198 | 29.44 | -0.942385 | 5.00E-05 | 0.00896 |
| ENSMUST00000064659 | ENSMUSG00000052676 | Zmat1 | 3501 | 2.53976 | 3.56726 | -0.490126 | 5.00E-05 | 0.00896 |
| ENSMUST00000113211 | ENSMUSG00000079435 | Rpl36a | 483 | 565.82 | 419.727 | 0.430891 | 5.00E-05 | 0.00896 |
| ENSMUST00000076038 | ENSMUSG00000038048 | Cntnap5c | 3918 | 2.27111 | 3.458 | -0.606537 | 5.00E-05 | 0.00896 |
| ENSMUST00000135562 | ENSMUSG00000030235 | Slco1c1 | 2680 | 9.55078 | 18.3024 | -0.938342 | 5.00E-05 | 0.00896 |
| ENSMUST00000102518 | ENSMUSG00000057530 | Ece1 | 4816 | 8.32631 | 5.85623 | 0.507705 | 5.00E-05 | 0.00896 |
| ENSMUST00000090781 | ENSMUSG00000068854 | Hist2h2be | 2620 | 2.97658 | 1.95352 | 0.607583 | 5.00E-05 | 0.00896 |
| ENSMUST00000090782 | ENSMUSG00000068855 | Hist2h2ac | 520 | 43.4868 | 15.9693 | 1.44528 | 5.00E-05 | 0.00896 |
| ENSMUST00000087342 | ENSMUSG00000067288 | Rps28 | 392 | 409.217 | 183.895 | 1.15398 | 5.00E-05 | 0.00896 |
| ENSMUST00000099703 | ENSMUSG00000075031 | Hist1h2bb | 452 | 42.8692 | 23.5081 | 0.866785 | 5.00E-05 | 0.00896 |
| ENSMUST00000075312 | ENSMUSG00000061808 | Ttr | 1219 | 226.713 | 147.489 | 0.620263 | 5.00E-05 | 0.00896 |
| ENSMUST00000001845 | ENSMUSG00000001794 | Capns1 | 1546 | 26.5548 | 1.99036 | 3.73787 | 5.00E-05 | 0.00896 |
| ENSMUST00000057885 | ENSMUSG00000047215 | Rpl9 | 738 | 957.42 | 748.437 | 0.355271 | 5.00E-05 | 0.00896 |
| ENSMUST00000106602 | ENSMUSG00000057322 | Rpl38 | 371 | 2596.25 | 1457.88 | 0.832554 | 5.00E-05 | 0.00896 |
| ENSMUST00000024706 | ENSMUSG00000023913 | Pla2g7 | 2097 | 60.0375 | 40.3662 | 0.572718 | 5.00E-05 | 0.00896 |
| ENSMUST00000178401 | ENSMUSG00000095325 | Zfp870 | 5126 | 2.97818 | 1.67623 | 0.829213 | 5.00E-05 | 0.00896 |
| ENSMUST00000057272 | ENSMUSG00000037270 | 4932438A13Rik | 15883 | 3.8288 | 20.4437 | -2.41669 | 5.00E-05 | 0.00896 |
| ENSMUST00000108995 | ENSMUSG00000089951 | Gm14435 | 643 | 12.1125 | 7.17816 | 0.754815 | 5.00E-05 | 0.00896 |
| ENSMUST00000084434 | ENSMUSG00000025508 | Rplp2 | 964 | 68.3132 | 43.1456 | 0.662951 | 5.00E-05 | 0.00896 |
| ENSMUST00000056549 | ENSMUSG00000047603 | Zfp235 | 3373 | 8.18825 | 5.23896 | 0.644276 | 5.00E-05 | 0.00896 |
| ENSMUST00000028199 | ENSMUSG00000026848 | Tor1b | 3039 | 3.77139 | 7.38624 | -0.969743 | 5.00E-05 | 0.00896 |
| ENSMUST00000153941 | ENSMUSG00000044712 | Slc38a6 | 3927 | 47.7476 | 80.042 | -0.745331 | 5.00E-05 | 0.00896 |
| ENSMUST00000028350 | ENSMUSG00000026974 | Zmynd19 | 3903 | 6.76252 | 3.21118 | 1.07446 | 5.00E-05 | 0.00896 |
| ENSMUST00000024811 | ENSMUSG00000024014 | Pim1 | 2638 | 2.32656 | 1.31343 | 0.824866 | 5.00E-05 | 0.00896 |
| ENSMUST00000191124 | ENSMUSG00000023826 | Park2 | 3202 | 1.91014 | 4.69131 | -1.29632 | 5.00E-05 | 0.00896 |
| ENSMUST00000020024 | ENSMUSG00000019874 | Fabp7 | 804 | 67.6485 | 31.663 | 1.09526 | 5.00E-05 | 0.00896 |
| ENSMUST00000076505 | ENSMUSG00000060224 | Pyroxd2 | 3119 | 0.279033 | 0.87761 | -1.65314 | 5.00E-05 | 0.00896 |
| ENSMUST00000010506 | ENSMUSG00000010362 | Rdm1 | 1124 | 4.55511 | 2.62492 | 0.795213 | 5.00E-05 | 0.00896 |
| ENSMUST00000029270 | ENSMUSG00000027715 | Ccna2 | 2965 | 0.445714 | 2.15425 | -2.273 | 5.00E-05 | 0.00896 |
| ENSMUST00000060792 | ENSMUSG00000004044 | Ptrf | 3491 | 4.1825 | 2.91016 | 0.523265 | 5.00E-05 | 0.00896 |
| ENSMUST00000192145 | ENSMUSG00000028080 | Lrba | 8843 | 2.17909 | 6.32824 | -1.53808 | 5.00E-05 | 0.00896 |
| ENSMUST00000135915 | ENSMUSG00000047617 | BC029214 | 896 | 4.34171 | 14.4895 | -1.73867 | 5.00E-05 | 0.00896 |
| ENSMUST00000006669 | ENSMUSG00000006494 | Pdk1 | 5185 | 20.2782 | 9.89082 | 1.03577 | 5.00E-05 | 0.00896 |
| ENSMUST00000102824 | ENSMUSG00000034459 | Ifit1 | 2656 | 1.33718 | 3.1783 | -1.24907 | 5.00E-05 | 0.00896 |
| ENSMUST00000102825 | ENSMUSG00000074896 | Ifit3 | 1745 | 5.01947 | 11.1151 | -1.14692 | 5.00E-05 | 0.00896 |
| ENSMUST00000116304 | ENSMUSG00000038642 | Ctss | 1355 | 54.1345 | 84.2908 | -0.638828 | 5.00E-05 | 0.00896 |
| ENSMUST00000164960 | ENSMUSG00000030134 | Rasgef1a | 3284 | 95.4798 | 69.075 | 0.467031 | 5.00E-05 | 0.00896 |
| ENSMUST00000165532 | ENSMUSG00000025794 | Rpl14 | 935 | 386.13 | 298.032 | 0.373617 | 5.00E-05 | 0.00896 |
| ENSMUST00000068045 | ENSMUSG00000054808 | Actn4 | 3877 | 10.1006 | 6.08247 | 0.731706 | 5.00E-05 | 0.00896 |
| ENSMUST00000062387 | ENSMUSG00000038026 | Kcnj9 | 3857 | 34.6862 | 24.6209 | 0.494482 | 5.00E-05 | 0.00896 |
| ENSMUST00000067230 | ENSMUSG00000076431 | Sox4 | 4795 | 6.33906 | 4.69718 | 0.432474 | 5.00E-05 | 0.00896 |
| ENSMUST00000063704 | ENSMUSG00000078496 | Gm13152 | 1244 | 0.661579 | 2.02632 | -1.61488 | 5.00E-05 | 0.00896 |
| ENSMUST00000073471 | ENSMUSG00000060938 | Rpl26 | 526 | 142.185 | 79.8311 | 0.832747 | 5.00E-05 | 0.00896 |
| ENSMUST00000068261 | ENSMUSG00000024403 | Atp6v1g2 | 1528 | 82.6508 | 118.232 | -0.516518 | 5.00E-05 | 0.00896 |
| ENSMUST00000109815 | ENSMUSG00000057897 | Camk2b | 3780 | 10.9686 | 34.3965 | -1.64888 | 5.00E-05 | 0.00896 |
| ENSMUST00000020203 | ENSMUSG00000020018 | Snrpf | 861 | 14.9396 | 9.87893 | 0.59671 | 5.00E-05 | 0.00896 |
| ENSMUST00000047309 | ENSMUSG00000035285 | Nat14 | 1612 | 34.4706 | 23.8487 | 0.53146 | 5.00E-05 | 0.00896 |

| | | | | | | | | |
|---|---|---|---|---|---|---|---|---|
| ENSMUST00000191829 | ENSMUSG00000102692 | Dchs2 | 10783 | 0.795445 | 1.32785 | -0.739261 | 5.00E-05 | 0.00896 |
| ENSMUST00000102852 | ENSMUSG00000050737 | Ptges | 3647 | 0.338866 | 1.20911 | -1.83516 | 5.00E-05 | 0.00896 |
| ENSMUST00000069803 | ENSMUSG00000089996 | Tmsb15b2 | 450 | 15.627 | 5.24189 | 1.57588 | 5.00E-05 | 0.00896 |
| ENSMUST00000053218 | ENSMUSG00000048826 | Dact2 | 2808 | 16.772 | 9.92186 | 0.757376 | 5.00E-05 | 0.00896 |
| ENSMUST00000102965 | ENSMUSG00000069266 | Hist1h4b | 397 | 127.671 | 36.6713 | 1.79971 | 5.00E-05 | 0.00896 |
| ENSMUST00000108205 | ENSMUSG00000059975 | Zfp74 | 3944 | 3.83711 | 2.23966 | 0.776741 | 5.00E-05 | 0.00896 |
| ENSMUST00000035269 | ENSMUSG00000032648 | Pygm | 2874 | 7.42257 | 5.12879 | 0.533299 | 5.00E-05 | 0.00896 |
| ENSMUST00000069562 | ENSMUSG00000055826 | Tescl | 855 | 0 | 0.491423 | #NAME? | 5.00E-05 | 0.00896 |
| ENSMUST00000002452 | ENSMUSG00000002379 | Ndufa11 | 2661 | 29.7707 | 19.3869 | 0.618811 | 5.00E-05 | 0.00896 |
| ENSMUST00000028259 | ENSMUSG00000026896 | Ifih1 | 5470 | 0.531701 | 1.25998 | -1.24471 | 5.00E-05 | 0.00896 |
| ENSMUST00000045604 | ENSMUSG00000036850 | Mrpl41 | 776 | 76.6941 | 42.9551 | 0.836288 | 5.00E-05 | 0.00896 |
| ENSMUST00000117242 | ENSMUSG00000034109 | Golim4 | 4551 | 2.85628 | 4.78613 | -0.744723 | 5.00E-05 | 0.00896 |
| ENSMUST00000087675 | ENSMUSG00000030225 | Dera | 1729 | 0.910446 | 2.084 | -1.19471 | 5.00E-05 | 0.00896 |
| ENSMUST00000026420 | ENSMUSG00000025362 | Rps26 | 447 | 623 | 348.463 | 0.838227 | 5.00E-05 | 0.00896 |
| ENSMUST00000105590 | ENSMUSG00000019768 | Esr1 | 6339 | 1.03891 | 0.637816 | 0.703859 | 5.00E-05 | 0.00896 |
| ENSMUST00000045356 | ENSMUSG00000041841 | Rpl37 | 867 | 612.114 | 434.803 | 0.493437 | 5.00E-05 | 0.00896 |
| ENSMUST00000079085 | ENSMUSG00000062006 | Rpl34 | 480 | 351.436 | 200.756 | 0.807818 | 5.00E-05 | 0.00896 |
| ENSMUST00000053458 | ENSMUSG00000050545 | Fam228b | 2886 | 5.98332 | 4.40112 | 0.443075 | 5.00E-05 | 0.00896 |
| ENSMUST00000050918 | ENSMUSG00000046447 | Camk2n1 | 4444 | 409.262 | 301.59 | 0.440433 | 5.00E-05 | 0.00896 |
| ENSMUST00000030765 | ENSMUSG00000028927 | Padi2 | 4774 | 3.29279 | 6.17897 | -0.908057 | 5.00E-05 | 0.00896 |
| ENSMUST00000065408 | ENSMUSG00000079018 | Ly6c1 | 844 | 34.4297 | 66.6129 | -0.952148 | 5.00E-05 | 0.00896 |
| ENSMUST00000077787 | ENSMUSG00000053985 | Zfp14 | 1506 | 8.45533 | 5.1465 | 0.716268 | 5.00E-05 | 0.00896 |
| ENSMUST00000174193 | ENSMUSG00000032517 | Mobp | 2609 | 144.755 | 217.097 | -0.584727 | 5.00E-05 | 0.00896 |
| ENSMUST00000025163 | ENSMUSG00000024309 | Pfdn6 | 807 | 77.8158 | 52.1912 | 0.576256 | 5.00E-05 | 0.00896 |
| ENSMUST00000062181 | ENSMUSG00000037029 | Zfp146 | 2024 | 22.7774 | 14.0409 | 0.697971 | 5.00E-05 | 0.00896 |
| ENSMUST00000123296 | ENSMUSG00000030869 | Ndufab1 | 1416 | 18.9984 | 10.349 | 0.876384 | 5.00E-05 | 0.00896 |
| ENSMUST00000080492 | ENSMUSG00000057863 | Rpl36 | 404 | 831.445 | 462.5 | 0.846167 | 5.00E-05 | 0.00896 |
| ENSMUST00000003416 | ENSMUSG00000048440 | Cyp4f16 | 2227 | 0.285478 | 0.834223 | -1.54705 | 5.00E-05 | 0.00896 |
| ENSMUST00000162772 | ENSMUSG00000030226 | Lmo3 | 3664 | 29.6863 | 17.9986 | 0.721911 | 5.00E-05 | 0.00896 |
| ENSMUST00000162170 | ENSMUSG00000030329 | Pianp | 2053 | 5.48344 | 10.2089 | -0.896678 | 5.00E-05 | 0.00896 |
| ENSMUST00000036370 | ENSMUSG00000033450 | Tagap | 3076 | 4.1425 | 2.4618 | 0.750787 | 5.00E-05 | 0.00896 |
| ENSMUST00000006618 | ENSMUSG00000028919 | Arhgef19 | 3414 | 0.783136 | 0.346706 | 1.17555 | 5.00E-05 | 0.00896 |
| ENSMUST00000172753 | ENSMUSG00000090877 | Hspa1b | 2803 | 5.44805 | 2.96057 | 0.879865 | 5.00E-05 | 0.00896 |
| ENSMUST00000140119 | ENSMUSG00000028626 | Col9a2 | 752 | 5.69885 | 1.9733 | 1.53006 | 5.00E-05 | 0.00896 |
| ENSMUST00000059980 | ENSMUSG00000046330 | Rpl37a | 584 | 1227.26 | 862.615 | 0.508656 | 5.00E-05 | 0.00896 |
| ENSMUST00000170122 | ENSMUSG00000090733 | Rps27 | 345 | 5085.51 | 2497.72 | 1.02578 | 5.00E-05 | 0.00896 |
| ENSMUST00000057486 | ENSMUSG00000048307 | Ankrd46 | 2480 | 76.1436 | 53.5047 | 0.509057 | 5.00E-05 | 0.00896 |
| ENSMUST00000034121 | ENSMUSG00000005142 | Man2b1 | 4325 | 5.17849 | 7.79428 | -0.589883 | 5.00E-05 | 0.00896 |
| ENSMUST00000043722 | ENSMUSG00000033880 | Lgals3bp | 2330 | 2.73436 | 4.20082 | -0.619466 | 5.00E-05 | 0.00896 |
| ENSMUST00000022239 | ENSMUSG00000021725 | Parp8 | 3088 | 7.81983 | 5.13526 | 0.6067 | 5.00E-05 | 0.00896 |
| ENSMUST00000034713 | ENSMUSG00000032193 | Ldlr | 4531 | 3.58401 | 5.0097 | -0.483149 | 5.00E-05 | 0.00896 |
| ENSMUST00000059650 | ENSMUSG00000043659 | Npsr1 | 3791 | 2.3859 | 1.00276 | 1.25057 | 5.00E-05 | 0.00896 |
| ENSMUST00000105813 | ENSMUSG00000041241 | Mul1 | 3944 | 9.2843 | 5.84903 | 0.666595 | 5.00E-05 | 0.00896 |
| ENSMUST00000035813 | ENSMUSG00000032827 | Ppp1r9a | 9547 | 19.7786 | 61.7864 | -1.64335 | 5.00E-05 | 0.00896 |
| ENSMUST00000206362 | ENSMUSG00000074283 | Zfp109 | 2932 | 2.59908 | 0.608645 | 2.09433 | 5.00E-05 | 0.00896 |
| ENSMUST00000093209 | ENSMUSG00000069919 | Hba-a1 | 699 | 44.6598 | 18.7312 | 1.25354 | 5.00E-05 | 0.00896 |
| ENSMUST00000030432 | ENSMUSG00000028672 | Hmgcl | 1418 | 8.22921 | 13.7653 | -0.742205 | 5.00E-05 | 0.00896 |
| ENSMUST00000030029 | ENSMUSG00000028344 | Invs | 5633 | 2.21109 | 5.77711 | -1.38559 | 5.00E-05 | 0.00896 |
| ENSMUST00000029649 | ENSMUSG00000028015 | Ctso | 3468 | 10.4101 | 6.01591 | 0.791133 | 5.00E-05 | 0.00896 |
| ENSMUST00000133544 | ENSMUSG00000026848 | Tor1b | 4775 | 2.84898 | 4.49202 | -0.656917 | 5.00E-05 | 0.00896 |
| ENSMUST00000098780 | ENSMUSG00000074358 | Cedc61 | 2022 | 2.03734 | 0.806223 | 1.33744 | 5.00E-05 | 0.00896 |
| ENSMUST00000035775 | ENSMUSG00000035215 | Lsm7 | 468 | 183.969 | 130.09 | 0.499949 | 5.00E-05 | 0.00896 |
| ENSMUST00000052867 | ENSMUSG00000050299 | Gm9843 | 504 | 22.1659 | 7.98555 | 1.47288 | 5.00E-05 | 0.00896 |
| ENSMUST00000052865 | ENSMUSG00000050912 | Tmem123 | 2907 | 4.1258 | 5.8852 | -0.512416 | 5.00E-05 | 0.00896 |
| ENSMUST00000028200 | ENSMUSG00000026849 | Tor1a | 1401 | 8.4235 | 12.7569 | -0.59879 | 5.00E-05 | 0.00896 |
| ENSMUST00000042035 | ENSMUSG00000034730 | Adgrb1 | 6228 | 11.3787 | 19.274 | -0.760319 | 5.00E-05 | 0.00896 |
| ENSMUST00000038108 | ENSMUSG00000037152 | Ndufc1 | 1249 | 25.2393 | 14.6949 | 0.780356 | 5.00E-05 | 0.00896 |
| ENSMUST00000102476 | ENSMUSG00000060802 | B2m | 860 | 64.6002 | 86.9981 | -0.429444 | 5.00E-05 | 0.00896 |
| ENSMUST00000041649 | ENSMUSG00000045027 | Prss22 | 1323 | 0.213449 | 3.77715 | -4.14533 | 5.00E-05 | 0.00896 |
| ENSMUST00000027343 | ENSMUSG00000026158 | Ogfrl1 | 4844 | 28.857 | 40.542 | -0.490495 | 5.00E-05 | 0.00896 |
| ENSMUST00000079251 | ENSMUSG00000058385 | Hist1h2bg | 477 | 71.6199 | 39.9155 | 0.843413 | 5.00E-05 | 0.00896 |
| ENSMUST00000150650 | ENSMUSG00000095362 | Gm14325 | 310 | 212.21 | 91.5017 | 1.21362 | 5.00E-05 | 0.00896 |
| ENSMUST00000076364 | ENSMUSG00000058443 | Rpl10-ps3 | 742 | 3.55259 | 1.30039 | 1.44992 | 5.00E-05 | 0.00896 |
| ENSMUST00000170872 | ENSMUSG00000023885 | Thbs2 | 5922 | 2.39406 | 3.53725 | -0.563166 | 5.00E-05 | 0.00896 |
| ENSMUST00000051386 | ENSMUSG00000038775 | Vill | 2909 | 1.22209 | 0.107622 | 3.50531 | 5.00E-05 | 0.00896 |
| ENSMUST00000076373 | ENSMUSG00000063696 | Gm8730 | 1075 | 9.0201 | 14.2366 | -0.658394 | 5.00E-05 | 0.00896 |
| ENSMUST00000078804 | ENSMUSG00000060636 | Rpl35a | 491 | 266.723 | 119.589 | 1.15726 | 5.00E-05 | 0.00896 |
| ENSMUST00000076941 | ENSMUSG00000033222 | Ttf2 | 4541 | 0.540211 | 0.217241 | 1.31423 | 5.00E-05 | 0.00896 |
| ENSMUST00000050433 | ENSMUSG00000031997 | Trpc6 | 3029 | 22.0241 | 4.3683 | 2.33394 | 5.00E-05 | 0.00896 |
| ENSMUST00000035105 | ENSMUSG00000032518 | Rpsa | 1861 | 261.72 | 106.416 | 1.2983 | 5.00E-05 | 0.00896 |
| ENSMUST00000051867 | ENSMUSG00000031683 | Lsm6 | 2107 | 12.2025 | 6.79629 | 0.84436 | 5.00E-05 | 0.00896 |
| ENSMUST00000042360 | ENSMUSG00000039676 | Capsl | 946 | 4.85154 | 2.15152 | 1.17308 | 5.00E-05 | 0.00896 |
| ENSMUST00000118012 | ENSMUSG00000078877 | Gm14295 | 1213 | 37.7303 | 27.0797 | 0.47851 | 5.00E-05 | 0.00896 |
| ENSMUST00000161741 | ENSMUSG00000038690 | Atp5j2 | 955 | 167.063 | 95.2071 | 0.811253 | 5.00E-05 | 0.00896 |
| ENSMUST00000092227 | ENSMUSG00000069539 | Scyl2 | 3566 | 6.27475 | 2.8119 | 1.15801 | 5.00E-05 | 0.00896 |
| ENSMUST00000042844 | ENSMUSG00000041120 | Nbl1 | 1793 | 23.6012 | 14.7838 | 0.674844 | 5.00E-05 | 0.00896 |
| ENSMUST00000028403 | ENSMUSG00000027015 | Cybrd1 | 5339 | 0.694289 | 0.119441 | 2.53923 | 5.00E-05 | 0.00896 |
| ENSMUST00000021719 | ENSMUSG00000021290 | 2010107E04Rik | 365 | 1704.12 | 1109.23 | 0.619465 | 5.00E-05 | 0.00896 |
| ENSMUST00000065678 | ENSMUSG00000053317 | Sec61b | 564 | 50.4096 | 21.7952 | 1.20969 | 5.00E-05 | 0.00896 |
| ENSMUST00000105106 | ENSMUSG00000069268 | Hist1h2bf | 461 | 13.8448 | 7.33599 | 0.916277 | 5.00E-05 | 0.00896 |

| | | | | | | | | |
|---|---|---|---|---|---|---|---|---|
| ENSMUST00000016396 | ENSMUSG00000016252 | Atp5e | 421 | 659.236 | 425.563 | 0.631425 | 5.00E-05 | 0.00896 |
| ENSMUST00000066070 | ENSMUSG00000053565 | Eif3k | 773 | 149.591 | 111.681 | 0.421644 | 5.00E-05 | 0.00896 |
| ENSMUST00000067556 | ENSMUSG00000044906 | 4930503L19Rik | 2291 | 1.81545 | 3.55097 | -0.967881 | 5.00E-05 | 0.00896 |
| ENSMUST00000032796 | ENSMUSG00000011427 | Zfp790 | 4525 | 6.94077 | 3.86694 | 0.843903 | 5.00E-05 | 0.00896 |
| ENSMUST00000159879 | ENSMUSG00000018199 | Trove2 | 8738 | 23.135 | 17.4794 | 0.40442 | 5.00E-05 | 0.00896 |
| ENSMUST00000148810 | ENSMUSG00000026483 | Fam129a | 6531 | 0.372165 | 0.682388 | -0.87465 | 5.00E-05 | 0.00896 |
| ENSMUST00000029722 | ENSMUSG00000028081 | Rps3a1 | 977 | 568.967 | 127.274 | 2.1604 | 5.00E-05 | 0.00896 |
| ENSMUST00000112640 | ENSMUSG00000026389 | Steap3 | 2752 | 1.08244 | 0.460961 | 1.23157 | 5.00E-05 | 0.00896 |
| ENSMUST00000102982 | ENSMUSG00000069308 | Hist1h2bp | 469 | 8.87623 | 4.03283 | 1.13816 | 5.00E-05 | 0.00896 |
| ENSMUST00000047687 | ENSMUSG00000041608 | Entpd3 | 3444 | 3.371 | 2.13224 | 0.660808 | 5.00E-05 | 0.00896 |
| ENSMUST00000113711 | ENSMUSG00000039715 | Wdr34 | 1814 | 18.698 | 11.8589 | 0.656909 | 5.00E-05 | 0.00896 |
| ENSMUST00000091756 | ENSMUSG00000094338 | Hist1h2bl | 381 | 61.2651 | 33.0844 | 0.888913 | 5.00E-05 | 0.00896 |
| ENSMUST00000048915 | ENSMUSG00000038374 | Rbm8a | 2608 | 10.5087 | 5.92786 | 0.825994 | 5.00E-05 | 0.00896 |
| ENSMUST00000183125 | ENSMUSG00000042063 | Zfp386 | 4779 | 5.26037 | 0.558041 | 3.23672 | 5.00E-05 | 0.00896 |
| ENSMUST00000030372 | ENSMUSG00000028626 | Col9a2 | 2924 | 2.36694 | 1.49336 | 0.664457 | 5.00E-05 | 0.00896 |
| ENSMUST00000028241 | ENSMUSG00000026880 | Stom | 2787 | 2.01639 | 4.99462 | -1.3086 | 5.00E-05 | 0.00896 |
| ENSMUST00000119103 | ENSMUSG00000032066 | Bco2 | 2118 | 0.858622 | 1.58397 | -0.883452 | 5.00E-05 | 0.00896 |
| ENSMUST00000045224 | ENSMUSG00000038007 | Acer2 | 4216 | 2.02277 | 4.52223 | -1.1607 | 5.00E-05 | 0.00896 |
| ENSMUST00000037134 | ENSMUSG00000040424 | Hipk4 | 2926 | 5.79172 | 8.54023 | -0.560282 | 5.00E-05 | 0.00896 |
| ENSMUST00000028239 | ENSMUSG00000026879 | Gsn | 2653 | 7.34721 | 13.095 | -0.833745 | 5.00E-05 | 0.00896 |
| ENSMUST00000020957 | ENSMUSG00000020629 | Adi1 | 1652 | 8.36156 | 4.77546 | 0.808132 | 5.00E-05 | 0.00896 |
| ENSMUST00000040907 | ENSMUSG00000036775 | Decr2 | 2187 | 20.4667 | 9.6629 | 1.08275 | 5.00E-05 | 0.00896 |
| ENSMUST00000015449 | ENSMUSG00000015305 | Sash1 | 7183 | 7.87588 | 5.85806 | 0.427017 | 5.00E-05 | 0.00896 |
| ENSMUST00000099294 | ENSMUSG00000025085 | Ablim1 | 6232 | 10.9835 | 17.1213 | -0.640452 | 5.00E-05 | 0.00896 |
| ENSMUST00000028332 | ENSMUSG00000026958 | Dpp7 | 1682 | 8.20839 | 13.3786 | -0.704755 | 5.00E-05 | 0.00896 |
| ENSMUST00000078369 | ENSMUSG00000061615 | Hist1h2ab | 473 | 12.5317 | 6.07962 | 1.04352 | 5.00E-05 | 0.00896 |
| ENSMUST00000015664 | ENSMUSG00000028111 | Ctsk | 1512 | 1.77449 | 3.06485 | -0.788415 | 5.00E-05 | 0.00896 |
| ENSMUST00000030851 | ENSMUSG00000028998 | Tomm7 | 1125 | 99.8208 | 75.0091 | 0.412275 | 5.00E-05 | 0.00896 |
| ENSMUST00000179734 | ENSMUSG00000095779 | Gm2163 | 2723 | 13.13 | 17.8309 | -0.441511 | 5.00E-05 | 0.00896 |
| ENSMUST00000102977 | ENSMUSG00000060639 | Hist1h4i | 727 | 105.731 | 68.5888 | 0.62435 | 5.00E-05 | 0.00896 |
| ENSMUST00000037820 | ENSMUSG00000038422 | Hdhd3 | 1205 | 9.46883 | 6.06538 | 0.642588 | 5.00E-05 | 0.00896 |
| ENSMUST00000102972 | ENSMUSG00000060981 | Hist1h4h | 473 | 652.241 | 397.655 | 0.713886 | 5.00E-05 | 0.00896 |
| ENSMUST00000027271 | ENSMUSG00000026102 | Inpp1 | 5767 | 3.21345 | 5.61024 | -0.803942 | 5.00E-05 | 0.00896 |
| ENSMUST00000102971 | ENSMUSG00000069274 | Hist1h4f | 391 | 279.013 | 189.5 | 0.558133 | 5.00E-05 | 0.00896 |
| ENSMUST00000029183 | ENSMUSG00000027654 | Fam83d | 2266 | 0.537131 | 1.2313 | -1.19684 | 5.00E-05 | 0.00896 |
| ENSMUST00000055071 | ENSMUSG00000079017 | Ifi27l2a | 463 | 3.57924 | 20.3631 | -2.50823 | 5.00E-05 | 0.00896 |
| ENSMUST00000102979 | ENSMUSG00000069305 | Hist1h4n | 430 | 407.973 | 216.772 | 0.912295 | 5.00E-05 | 0.00896 |
| ENSMUST00000042665 | ENSMUSG00000034422 | Parp14 | 7417 | 0.352579 | 0.603346 | -0.775039 | 5.00E-05 | 0.00896 |
| ENSMUST00000135776 | ENSMUSG00000046079 | Lrrc8d | 5736 | 1.49413 | 3.55164 | -1.24918 | 5.00E-05 | 0.00896 |
| ENSMUST00000194649 | ENSMUSG00000039221 | Rpl22l1 | 525 | 183.389 | 14.9086 | 3.62069 | 5.00E-05 | 0.00896 |
| ENSMUST00000025572 | ENSMUSG00000024670 | Cd6 | 2173 | 0.414124 | 1.13289 | -1.45187 | 5.00E-05 | 0.00896 |
| ENSMUST00000113599 | ENSMUSG00000056492 | Adgrf5 | 8535 | 2.37509 | 3.52314 | -0.56888 | 5.00E-05 | 0.00896 |
| ENSMUST00000205671 | ENSMUSG00000030604 | Zfp626 | 5329 | 5.21657 | 3.86224 | 0.433664 | 5.00E-05 | 0.00896 |
| ENSMUST00000127027 | ENSMUSG00000096696 | Zfp960 | 2667 | 1.21942 | 2.16247 | -0.826482 | 5.00E-05 | 0.00896 |
| ENSMUST00000009039 | ENSMUSG00000058600 | Rpl30 | 570 | 212.942 | 128.633 | 0.727199 | 5.00E-05 | 0.00896 |
| ENSMUST00000022751 | ENSMUSG00000022151 | Ttc33 | 1832 | 36.6084 | 17.3615 | 1.07629 | 5.00E-05 | 0.00896 |
| ENSMUST00000187028 | ENSMUSG00000099689 | Zfp383 | 3208 | 3.20396 | 1.4682 | 1.12581 | 5.00E-05 | 0.00896 |
| ENSMUST00000030538 | ENSMUSG00000028757 | Ddost | 2120 | 46.4563 | 34.9394 | 0.411021 | 5.00E-05 | 0.00896 |
| ENSMUST00000048718 | ENSMUSG00000037022 | Mmaa | 2843 | 7.73067 | 5.73656 | 0.430409 | 5.00E-05 | 0.00896 |
| ENSMUST00000142555 | ENSMUSG00000069919 | Hba-a1 | 494 | 249.843 | 143.494 | 0.800036 | 5.00E-05 | 0.00896 |
| ENSMUST00000089430 | ENSMUSG00000068240 | Gm11808 | 504 | 8.4874 | 3.8649 | 1.13489 | 5.00E-05 | 0.00896 |
| ENSMUST00000065103 | ENSMUSG00000052962 | Mrpl35 | 3721 | 5.92122 | 1.78235 | 1.73211 | 5.00E-05 | 0.00896 |
| ENSMUST00000097992 | ENSMUSG00000038368 | Focad | 5791 | 10.508 | 17.0206 | -0.695788 | 5.00E-05 | 0.00896 |
| ENSMUST00000118193 | ENSMUSG00000022151 | Ttc33 | 1671 | 17.8915 | 37.3801 | -1.06299 | 5.00E-05 | 0.00896 |
| ENSMUST00000167323 | ENSMUSG00000090698 | Apold1 | 3567 | 1.67313 | 2.74567 | -0.714606 | 5.00E-05 | 0.00896 |
| ENSMUST00000061852 | ENSMUSG00000026648 | Dclre1c | 7075 | 0.227636 | 0.887767 | -1.96345 | 5.00E-05 | 0.00896 |
| ENSMUST00000023934 | ENSMUSG00000052305 | Hbb-bs | 778 | 139.116 | 44.4335 | 1.64657 | 5.00E-05 | 0.00896 |
| ENSMUST00000063976 | ENSMUSG00000052214 | Opa3 | 3366 | 17.54 | 12.2107 | 0.522506 | 5.00E-05 | 0.00896 |
| ENSMUST00000078034 | ENSMUSG00000062456 | Rpl9-ps6 | 691 | 12.8409 | 4.25318 | 1.59413 | 5.00E-05 | 0.00896 |
| ENSMUST00000087605 | ENSMUSG00000024300 | Myo1f | 3819 | 0.610911 | 1.05402 | -0.786869 | 5.00E-05 | 0.00896 |
| ENSMUST00000046285 | ENSMUSG00000036887 | C1qa | 1040 | 35.8171 | 55.1394 | -0.622437 | 5.00E-05 | 0.00896 |
| ENSMUST00000053020 | ENSMUSG00000034413 | Neurl1b | 6195 | 11.0395 | 7.73735 | 0.512759 | 5.00E-05 | 0.00896 |
| ENSMUST00000066708 | ENSMUSG00000029307 | Dmp1 | 2764 | 0.366427 | 0.94718 | -1.37011 | 5.00E-05 | 0.00896 |
| ENSMUST00000108429 | ENSMUSG00000040952 | Rps19 | 633 | 319.416 | 167.407 | 0.932082 | 5.00E-05 | 0.00896 |
| ENSMUST00000028470 | ENSMUSG00000027076 | Timm10 | 690 | 40.2998 | 29.5209 | 0.449035 | 5.00E-05 | 0.00896 |
| ENSMUST00000030401 | ENSMUSG00000028648 | Ndufs5 | 525 | 294.022 | 224.207 | 0.391092 | 5.00E-05 | 0.00896 |
| ENSMUST00000039205 | ENSMUSG00000035473 | Galm | 2314 | 0.717184 | 1.48842 | -1.05337 | 5.00E-05 | 0.00896 |
| ENSMUST00000151224 | ENSMUSG00000009216 | Fam163b | 2959 | 9.44443 | 13.1439 | -0.47686 | 5.00E-05 | 0.00896 |
| ENSMUST00000165033 | ENSMUSG00000038418 | Egr1 | 3036 | 26.2421 | 39.8109 | -0.60128 | 5.00E-05 | 0.00896 |
| ENSMUST00000109626 | ENSMUSG00000036712 | Cyld | 7945 | 4.02991 | 13.7088 | -1.76628 | 5.00E-05 | 0.00896 |
| ENSMUST00000186475 | ENSMUSG00000099689 | Zfp383 | 2614 | 2.11192 | 1.00307 | 1.07413 | 5.00E-05 | 0.00896 |
| ENSMUST00000069507 | ENSMUSG00000073418 | C4b | 5379 | 5.28792 | 3.60957 | 0.550876 | 5.00E-05 | 0.00896 |
| ENSMUST00000040824 | ENSMUSG00000042737 | Dpm3 | 404 | 114.641 | 74.8697 | 0.614674 | 5.00E-05 | 0.00896 |
| ENSMUST00000058210 | ENSMUSG00000015766 | Eps8 | 4567 | 2.56078 | 6.32752 | -1.30505 | 5.00E-05 | 0.00896 |
| ENSMUST00000164044 | ENSMUSG00000059498 | Fcgr3 | 1347 | 5.6977 | 9.04075 | -0.666063 | 5.00E-05 | 0.00896 |
| ENSMUST00000114020 | ENSMUSG00000052406 | Rexo4 | 2278 | 19.2931 | 13.9905 | 0.463635 | 5.00E-05 | 0.00896 |
| ENSMUST00000069486 | ENSMUSG00000055760 | Gemin6 | 1134 | 5.91072 | 3.52901 | 0.744069 | 5.00E-05 | 0.00896 |
| ENSMUST00000006397 | ENSMUSG00000006235 | Epor | 1829 | 2.92082 | 1.70144 | 0.779614 | 5.00E-05 | 0.00896 |
| ENSMUST00000167969 | ENSMUSG00000090963 | Gm17655 | 255 | 78.2862 | 23.3409 | 1.74589 | 5.00E-05 | 0.00896 |
| ENSMUST00000024832 | ENSMUSG00000024033 | Rsph1 | 1164 | 1.5924 | 3.99338 | -1.32641 | 5.00E-05 | 0.00896 |

| | | | | | | | | |
|---|---|---|---|---|---|---|---|---|
| ENSMUST00000055738 | ENSMUSG00000031431 | Tsc22d3 | 1974 | 25.3661 | 42.3212 | -0.73848 | 5.00E-05 | 0.00896 |
| ENSMUST00000074455 | ENSMUSG00000066838 | Zfp772 | 2772 | 2.32771 | 4.6155 | -0.987576 | 5.00E-05 | 0.00896 |
| ENSMUST00000027931 | ENSMUSG00000026622 | Nek2 | 3227 | 0.550049 | 0.110751 | 2.31224 | 5.00E-05 | 0.00896 |
| ENSMUST00000024884 | ENSMUSG00000024079 | Eif2ak2 | 4313 | 2.03069 | 3.02105 | -0.573084 | 5.00E-05 | 0.00896 |
| ENSMUST00000104998 | ENSMUSG00000078201 | Tmem203 | 854 | 9.4241 | 17.9501 | -0.929565 | 5.00E-05 | 0.00896 |
| ENSMUST00000051672 | ENSMUSG00000046718 | Bst2 | 764 | 1.21302 | 3.73873 | -1.62395 | 5.00E-05 | 0.00896 |
| ENSMUST00000082411 | ENSMUSG00000064360 | mt-Nd3 | 348 | 2470.25 | 1085.54 | 1.18624 | 5.00E-05 | 0.00896 |
| ENSMUST00000061688 | ENSMUSG00000036552 | Ermard | 2120 | 5.91017 | 2.13401 | 1.46963 | 5.00E-05 | 0.00896 |
| ENSMUST00000086006 | ENSMUSG00000087598 | Zfp111 | 6402 | 1.57485 | 0.85891 | 0.874635 | 5.00E-05 | 0.00896 |
| ENSMUST00000032696 | ENSMUSG00000055305 | Zfp93 | 2324 | 4.58769 | 1.91569 | 1.2599 | 5.00E-05 | 0.00896 |
| ENSMUST00000076249 | ENSMUSG00000062488 | Ifit3b | 2000 | 1.94827 | 3.80832 | -0.966963 | 5.00E-05 | 0.00896 |
| ENSMUST00000090287 | ENSMUSG00000018830 | Myh11 | 6632 | 1.64878 | 2.55992 | -0.634703 | 5.00E-05 | 0.00896 |
| ENSMUST00000111518 | ENSMUSG00000040181 | Fmo1 | 1506 | 8.70814 | 29.6668 | -1.76841 | 5.00E-05 | 0.00896 |
| ENSMUST00000144826 | ENSMUSG00000023087 | Noct | 9797 | 19.8229 | 34.4388 | -0.796866 | 5.00E-05 | 0.00896 |
| ENSMUST00000118009 | ENSMUSG00000014907 | Naf1 | 2990 | 3.71575 | 2.35546 | 0.657644 | 5.00E-05 | 0.00896 |
| ENSMUST00000171588 | ENSMUSG00000054263 | Lifr | 9655 | 10.7193 | 6.19771 | 0.790399 | 5.00E-05 | 0.00896 |
| ENSMUST00000103146 | ENSMUSG00000071415 | Rpl23 | 1059 | 269.016 | 183.122 | 0.554883 | 5.00E-05 | 0.00896 |
| ENSMUST00000179285 | ENSMUSG00000096010 | Hist4h4 | 2863 | 25.7768 | 19.0595 | 0.435559 | 5.00E-05 | 0.00896 |
| ENSMUST00000090746 | ENSMUSG00000027875 | Hmgcs2 | 3290 | 3.94851 | 2.58729 | 0.609868 | 5.00E-05 | 0.00896 |
| ENSMUST00000066489 | ENSMUSG00000006931 | P3h4 | 2178 | 8.39034 | 4.12607 | 1.02396 | 5.00E-05 | 0.00896 |
| ENSMUST00000082396 | ENSMUSG00000064345 | mt-Nd2 | 1038 | 563.843 | 435.064 | 0.374067 | 5.00E-05 | 0.00896 |
| ENSMUST00000179505 | ENSMUSG00000095041 | PISD | 2373 | 39.8585 | 9.28562 | 2.10182 | 5.00E-05 | 0.00896 |
| ENSMUST00000045756 | ENSMUSG00000041959 | S100a10 | 685 | 48.5343 | 31.6277 | 0.617816 | 5.00E-05 | 0.00896 |
| ENSMUST00000055296 | ENSMUSG00000049553 | Polr1a | 9216 | 2.17232 | 0.869029 | 1.32176 | 5.00E-05 | 0.00896 |
| ENSMUST00000150108 | ENSMUSG00000013593 | Ndufs2 | 662 | 130.327 | 8.94722 | 3.86455 | 5.00E-05 | 0.00896 |
| ENSMUST00000043870 | ENSMUSG00000038489 | Polr2l | 1650 | 24.5994 | 17.0954 | 0.525012 | 5.00E-05 | 0.00896 |
| ENSMUST00000063683 | ENSMUSG00000052031 | Tagap1 | 2358 | 6.89182 | 13.2426 | -0.942233 | 5.00E-05 | 0.00896 |
| ENSMUST00000023994 | ENSMUSG00000023224 | Serping1 | 1768 | 1.65432 | 2.74343 | -0.729742 | 5.00E-05 | 0.00896 |
| ENSMUST00000044681 | ENSMUSG00000035199 | Arl6ip5 | 1442 | 85.995 | 60.9559 | 0.496488 | 5.00E-05 | 0.00896 |
| ENSMUST00000187126 | ENSMUSG00000093803 | Ppp2r3d | 254 | 94.1352 | 25.4388 | 1.8877 | 5.00E-05 | 0.00896 |
| ENSMUST00000098822 | ENSMUSG00000030386 | Zfp606 | 4748 | 4.02061 | 2.30501 | 0.802641 | 5.00E-05 | 0.00896 |
| ENSMUST00000180180 | ENSMUSG00000066839 | Ecsit | 1676 | 3.55188 | 10.5888 | -1.57588 | 5.00E-05 | 0.00896 |
| ENSMUST00000113071 | ENSMUSG00000089768 | Tmsb15b1 | 439 | 12.4684 | 3.05487 | 2.02909 | 5.00E-05 | 0.00896 |
| ENSMUST00000020145 | ENSMUSG00000019970 | Sgk1 | 2456 | 26.657 | 44.8449 | -0.750429 | 5.00E-05 | 0.00896 |
| ENSMUST00000075602 | ENSMUSG00000070713 | Gm10282 | 1159 | 1.01069 | 3.69247 | -1.86924 | 5.00E-05 | 0.00896 |
| ENSMUST00000048248 | ENSMUSG00000035390 | Brsk1 | 2996 | 19.5244 | 33.3767 | -0.77356 | 5.00E-05 | 0.00896 |
| ENSMUST00000029846 | ENSMUSG00000028195 | Cyr61 | 2019 | 2.24778 | 3.91789 | -0.801575 | 5.00E-05 | 0.00896 |
| ENSMUST00000023207 | ENSMUSG00000022548 | Apod | 979 | 11.5188 | 36.9872 | -1.68303 | 5.00E-05 | 0.00896 |
| ENSMUST00000047321 | ENSMUSG00000055116 | Arntl | 2904 | 17.866 | 13.4877 | 0.405572 | 5.00E-05 | 0.00896 |
| ENSMUST00000087328 | ENSMUSG00000091971 | Hspa1a | 2967 | 3.63064 | 2.10996 | 0.783007 | 5.00E-05 | 0.00896 |
| ENSMUST00000002176 | ENSMUSG00000002107 | Celf2 | 7718 | 14.2986 | 4.17747 | 1.77517 | 5.00E-05 | 0.00896 |
| ENSMUST00000105285 | ENSMUSG00000019936 | Epyc | 2539 | 0.507597 | 1.14033 | -1.1677 | 5.00E-05 | 0.00896 |
| ENSMUST00000138999 | ENSMUSG00000026568 | Mpc2 | 493 | 79.5992 | 42.867 | 0.892884 | 5.00E-05 | 0.00896 |
| ENSMUST00000094934 | ENSMUSG00000070880 | Gad1 | 3355 | 94.1115 | 63.3583 | 0.570837 | 5.00E-05 | 0.00896 |
| ENSMUST00000096014 | ENSMUSG00000071528 | Usmg5 | 341 | 1446.42 | 979.181 | 0.562836 | 5.00E-05 | 0.00896 |
| ENSMUST00000081134 | ENSMUSG00000060445 | Sycp2 | 5717 | 0.49954 | 1.06219 | -1.08838 | 5.00E-05 | 0.00896 |
| ENSMUST00000064061 | ENSMUSG00000060257 | Scrt2 | 3395 | 0.911096 | 1.46403 | -0.684265 | 5.00E-05 | 0.00896 |
| ENSMUST00000075304 | ENSMUSG00000041488 | Stx3 | 2457 | 3.44633 | 10.1946 | -1.56467 | 5.00E-05 | 0.00896 |
| ENSMUST00000106635 | ENSMUSG00000051497 | Kcnj16 | 3671 | 1.3313 | 4.60352 | -1.7899 | 5.00E-05 | 0.00896 |
| ENSMUST00000179436 | ENSMUSG00000095742 | CAAA01147332.1 | 251 | 93.4701 | 35.7207 | 1.38774 | 5.00E-05 | 0.00896 |
| ENSMUST00000034881 | ENSMUSG00000032330 | Cox7a2 | 754 | 387.097 | 273.286 | 0.502285 | 5.00E-05 | 0.00896 |
| ENSMUST00000167994 | ENSMUSG00000051977 | Prdm9 | 3460 | 0.185171 | 0.645088 | -1.80064 | 5.00E-05 | 0.00896 |
| ENSMUST00000028187 | ENSMUSG00000026840 | Lamc3 | 5871 | 0.542745 | 1.00551 | -0.889574 | 5.00E-05 | 0.00896 |
| ENSMUST00000046223 | ENSMUSG00000041828 | Abca8a | 5614 | 3.01258 | 5.34578 | -0.8274 | 5.00E-05 | 0.00896 |
| ENSMUST00000172815 | ENSMUSG00000092216 | Gm19345 | 998 | 1.85128 | 0.458993 | 2.01198 | 5.00E-05 | 0.00896 |
| ENSMUST00000112186 | ENSMUSG00000041975 | Mettl8 | 2347 | 3.46499 | 5.90488 | -0.769055 | 5.00E-05 | 0.00896 |
| ENSMUST00000062765 | ENSMUSG00000030443 | Zfp583 | 2550 | 1.09412 | 2.31636 | -1.08209 | 5.00E-05 | 0.00896 |
| ENSMUST00000059080 | ENSMUSG00000039001 | Rps21 | 394 | 2945.25 | 1783.1 | 0.724003 | 5.00E-05 | 0.00896 |
| ENSMUST00000051364 | ENSMUSG00000044709 | Gemin7 | 763 | 18.1975 | 8.48316 | 1.10106 | 5.00E-05 | 0.00896 |
| ENSMUST00000102964 | ENSMUSG00000060093 | Hist1h4a | 539 | 202.001 | 130.893 | 0.625971 | 5.00E-05 | 0.00896 |
| ENSMUST00000102967 | ENSMUSG00000060678 | Hist1h4c | 395 | 765.816 | 275.503 | 1.47493 | 5.00E-05 | 0.00896 |
| ENSMUST00000060716 | ENSMUSG00000051354 | Samd3 | 1874 | 0.267962 | 0.949173 | -1.82464 | 5.00E-05 | 0.00896 |
| ENSMUST00000102969 | ENSMUSG00000069272 | Hist1h2ae | 460 | 17.1223 | 10.0728 | 0.765405 | 5.00E-05 | 0.00896 |
| ENSMUST00000102968 | ENSMUSG00000061482 | Hist1h4d | 393 | 837.585 | 510.133 | 0.715363 | 5.00E-05 | 0.00896 |
| ENSMUST00000028764 | ENSMUSG00000027301 | Oxt | 537 | 38.6873 | 21.5946 | 0.841187 | 5.00E-05 | 0.00896 |
| ENSMUST00000029269 | ENSMUSG00000027714 | Exosc9 | 1617 | 14.24 | 21.9267 | -0.622736 | 5.00E-05 | 0.00896 |
| ENSMUST00000028045 | ENSMUSG00000026712 | Mrc1 | 5374 | 1.62197 | 2.53453 | -0.643968 | 5.00E-05 | 0.00896 |
| ENSMUST00000029266 | ENSMUSG00000027712 | Anxa5 | 1731 | 59.5502 | 34.8154 | 0.774383 | 5.00E-05 | 0.00896 |
| ENSMUST00000070631 | ENSMUSG00000037921 | Ddx60 | 5992 | 0.101225 | 0.447987 | -2.14589 | 5.00E-05 | 0.00896 |
| ENSMUST00000180159 | ENSMUSG00000045996 | Polr2k | 489 | 59.3927 | 39.0327 | 0.605601 | 5.00E-05 | 0.00896 |
| ENSMUST00000086535 | ENSMUSG00000073702 | Rpl31 | 1049 | 219.21 | 142.404 | 0.622319 | 5.00E-05 | 0.00896 |
| ENSMUST00000038423 | ENSMUSG00000033355 | Rtp4 | 1573 | 0.791804 | 1.72829 | -1.12613 | 5.00E-05 | 0.00896 |
| ENSMUST00000020102 | ENSMUSG00000019935 | Slc17a8 | 4456 | 0.578833 | 1.62741 | -1.49136 | 5.00E-05 | 0.00896 |
| ENSMUST00000049628 | ENSMUSG00000050856 | Atp5k | 371 | 842.961 | 601.646 | 0.486551 | 5.00E-05 | 0.00896 |
| ENSMUST00000166950 | ENSMUSG00000078974 | Sec61g | 754 | 84.2193 | 50.9388 | 0.725386 | 5.00E-05 | 0.00896 |
| ENSMUST00000054544 | ENSMUSG00000049751 | Rpl36al | 526 | 270.383 | 189.183 | 0.515221 | 5.00E-05 | 0.00896 |
| ENSMUST00000127404 | ENSMUSG00000072955 | Tmsb15l | 816 | 6.5897 | 1.58324 | 2.05733 | 5.00E-05 | 0.00896 |
| ENSMUST00000043938 | ENSMUSG00000038910 | Plcl2 | 4050 | 49.1632 | 66.9912 | -0.446394 | 5.00E-05 | 0.00896 |
| ENSMUST00000025025 | ENSMUSG00000024190 | Dusp1 | 1990 | 9.07346 | 16.5291 | -0.865282 | 5.00E-05 | 0.00896 |
| ENSMUST00000029406 | ENSMUSG00000027824 | Vmn2r1 | 2739 | 0.157774 | 0.650253 | -2.04314 | 5.00E-05 | 0.00896 |

| | | | | | | | | |
|---|---|---|---|---|---|---|---|---|
| ENSMUST00000029568 | ENSMUSG00000027956 | Tmem144 | 2898 | 4.83194 | 3.1073 | 0.636939 | 5.00E-05 | 0.00896 |
| ENSMUST00000087176 | ENSMUSG00000050121 | Opalin | 1838 | 4.63874 | 6.81319 | -0.554597 | 5.00E-05 | 0.00896 |
| ENSMUST00000001051 | ENSMUSG00000001025 | S100a6 | 731 | 24.7133 | 15.9178 | 0.634647 | 5.00E-05 | 0.00896 |
| ENSMUST00000058030 | ENSMUSG00000062937 | Mtap | 2797 | 4.50295 | 2.49181 | 0.853675 | 5.00E-05 | 0.00896 |
| ENSMUST00000021674 | ENSMUSG00000021250 | Fos | 2108 | 11.3983 | 17.494 | -0.618047 | 5.00E-05 | 0.00896 |
| ENSMUST00000156844 | ENSMUSG00000078875 | Gm14419 | 752 | 17.263 | 10.1257 | 0.769661 | 5.00E-05 | 0.00896 |
| ENSMUST00000185187 | ENSMUSG00000061762 | Tac1 | 1230 | 29.5171 | 50.5625 | -0.776519 | 5.00E-05 | 0.00896 |
| ENSMUST00000133324 | ENSMUSG00000020672 | Sntg2 | 1830 | 3.06355 | 1.48267 | 1.047 | 5.00E-05 | 0.00896 |
| ENSMUST00000032279 | ENSMUSG00000030172 | Erc1 | 8356 | 0.674057 | 3.16908 | -2.23312 | 5.00E-05 | 0.00896 |
| ENSMUST00000164491 | ENSMUSG00000090667 | Gm765 | 3132 | 3.38658 | 2.21533 | 0.612311 | 5.00E-05 | 0.00896 |
| ENSMUST00000072271 | ENSMUSG00000019232 | Etnppl | 4468 | 1.90862 | 3.04858 | -0.67561 | 5.00E-05 | 0.00896 |
| ENSMUST00000100841 | ENSMUSG00000015766 | Eps8 | 4151 | 5.89444 | 1.35026 | 2.12612 | 5.00E-05 | 0.00896 |
| ENSMUST00000172785 | ENSMUSG00000073411 | H2-D1 | 2102 | 2.57622 | 4.23092 | -0.715718 | 5.00E-05 | 0.00896 |
| ENSMUST00000005077 | ENSMUSG00000004951 | Hspb1 | 903 | 9.16685 | 5.16758 | 0.826936 | 5.00E-05 | 0.00896 |
| ENSMUST00000085668 | ENSMUSG00000066647 | Gm5113 | 3002 | 8.30675 | 5.16282 | 0.686125 | 5.00E-05 | 0.00896 |
| ENSMUST00000178343 | ENSMUSG00000095041 | PISD | 4060 | 31.1831 | 11.4406 | 1.4466 | 5.00E-05 | 0.00896 |
| ENSMUST00000063492 | ENSMUSG00000051920 | Rspo2 | 3324 | 8.12312 | 3.80657 | 1.09354 | 5.00E-05 | 0.00896 |
| ENSMUST00000081982 | ENSMUSG00000037259 | Dzank1 | 6847 | 49.092 | 36.5921 | 0.423956 | 5.00E-05 | 0.00896 |
| ENSMUST00000068975 | ENSMUSG00000057101 | Zfp180 | 3901 | 24.1974 | 14.5214 | 0.736672 | 5.00E-05 | 0.00896 |
| ENSMUST00000103071 | ENSMUSG00000020713 | Gh | 862 | 3.01122 | 0.778706 | 1.95119 | 5.00E-05 | 0.00896 |
| ENSMUST00000135737 | ENSMUSG00000026839 | Upp2 | 2848 | 1.24781 | 7.08122 | -2.5046 | 5.00E-05 | 0.00896 |
| ENSMUST00000092639 | ENSMUSG00000019899 | Lama2 | 9614 | 1.86373 | 2.79903 | -0.586738 | 5.00E-05 | 0.00896 |
| ENSMUST00000006254 | ENSMUSG00000006095 | Tbcb | 1416 | 43.351 | 28.9989 | 0.580063 | 5.00E-05 | 0.00896 |
| ENSMUST00000109761 | ENSMUSG00000003813 | Rad23a | 1842 | 12.0066 | 5.20599 | 1.20559 | 5.00E-05 | 0.00896 |
| ENSMUST00000015236 | ENSMUSG00000015092 | Edf1 | 722 | 110.77 | 67.2698 | 0.719534 | 5.00E-05 | 0.00896 |
| ENSMUST00000065291 | ENSMUSG00000053070 | 9230110C19Rik | 1592 | 6.20737 | 9.22004 | -0.57079 | 5.00E-05 | 0.00896 |
| ENSMUST00000030317 | ENSMUSG00000028583 | Pdpn | 1817 | 6.97095 | 4.42714 | 0.654981 | 5.00E-05 | 0.00896 |
| ENSMUST00000094964 | ENSMUSG00000096141 | Dnah7a | 12245 | 0.550725 | 0.0589681 | 3.22333 | 5.00E-05 | 0.00896 |
| ENSMUST00000077687 | ENSMUSG00000036641 | Ccdc148 | 3977 | 5.19692 | 8.69583 | -0.742667 | 5.00E-05 | 0.00896 |
| ENSMUST00000051065 | ENSMUSG00000045441 | Gprin3 | 8339 | 8.81993 | 3.73444 | 1.23987 | 5.00E-05 | 0.00896 |
| ENSMUST00000035120 | ENSMUSG00000032532 | Cck | 685 | 464.126 | 357.779 | 0.37545 | 0.0001 | 0.01618 |
| ENSMUST00000195247 | ENSMUSG00000058407 | Txndc9 | 1699 | 4.41098 | 11.8596 | -1.42688 | 0.0001 | 0.01618 |
| ENSMUST00000090180 | ENSMUSG00000021904 | Sema3g | 4538 | 1.08574 | 1.64007 | -0.595077 | 0.0001 | 0.01618 |
| ENSMUST00000106956 | ENSMUSG00000061086 | Myl4 | 900 | 12.644 | 19.1987 | -0.602548 | 0.0001 | 0.01618 |
| ENSMUST00000162031 | ENSMUSG00000058407 | Txndc9 | 3381 | 9.93987 | 6.42442 | 0.62966 | 0.0001 | 0.01618 |
| ENSMUST00000179077 | ENSMUSG00000096768 | Erdr1 | 895 | 4.39688 | 1.68781 | 1.38133 | 0.0001 | 0.01618 |
| ENSMUST00000046807 | ENSMUSG00000037762 | Slc16a9 | 3756 | 0.436881 | 0.867582 | -0.989761 | 0.0001 | 0.01618 |
| ENSMUST00000189707 | ENSMUSG00000093803 | Ppp2r3d | 385 | 19.829 | 11.5103 | 0.784688 | 0.0001 | 0.01618 |
| ENSMUST00000084207 | ENSMUSG00000071176 | Arhgef10 | 5528 | 3.28917 | 4.74937 | -0.530014 | 0.0001 | 0.01618 |
| ENSMUST00000004203 | ENSMUSG00000004100 | Ppan | 1670 | 4.43374 | 6.67552 | -0.590356 | 0.0001 | 0.01618 |
| ENSMUST00000020768 | ENSMUSG00000020475 | Pgam2 | 840 | 5.35828 | 8.33572 | -0.637536 | 0.0001 | 0.01618 |
| ENSMUST00000105031 | ENSMUSG00000078234 | Klhdc7a | 7807 | 1.84315 | 1.31639 | 0.485585 | 0.0001 | 0.01618 |
| ENSMUST00000023538 | ENSMUSG00000022836 | Mylk | 7824 | 2.19574 | 2.98016 | -0.440686 | 0.0001 | 0.01618 |
| ENSMUST00000105365 | ENSMUSG00000045193 | Cirbp | 1321 | 42.6346 | 56.11 | -0.396232 | 0.0001 | 0.01618 |
| ENSMUST00000021114 | ENSMUSG00000020766 | Galk1 | 1406 | 6.07765 | 4.04111 | 0.588762 | 0.0001 | 0.01618 |
| ENSMUST00000089257 | ENSMUSG00000068154 | Insm1 | 3100 | 2.38484 | 1.59509 | 0.580251 | 0.0001 | 0.01618 |
| ENSMUST00000190422 | ENSMUSG00000101523 | Gm10031 | 2653 | 0.496325 | 0.165457 | 1.58482 | 0.0001 | 0.01618 |
| ENSMUST00000043867 | ENSMUSG00000039221 | Rpl22l1 | 552 | 61.8604 | 110.354 | -0.835057 | 0.0001 | 0.01618 |
| ENSMUST00000029331 | ENSMUSG00000027765 | P2ry1 | 3886 | 2.10826 | 0.966153 | 1.12573 | 0.0001 | 0.01618 |
| ENSMUST00000045537 | ENSMUSG00000040495 | Chrm4 | 1440 | 17.3709 | 12.3274 | 0.494798 | 0.0001 | 0.01618 |
| ENSMUST00000139045 | ENSMUSG00000041921 | Metap1d | 2770 | 2.79589 | 0.579358 | 2.27078 | 0.0001 | 0.01618 |
| ENSMUST00000023453 | ENSMUSG00000022769 | Sdf2l1 | 1108 | 18.9038 | 13.4144 | 0.494893 | 0.0001 | 0.01618 |
| ENSMUST00000111194 | ENSMUSG00000005087 | Cd44 | 5045 | 0.457981 | 1.02294 | -1.15936 | 0.0001 | 0.01618 |
| ENSMUST00000073080 | ENSMUSG00000058927 | Gm10053 | 949 | 1.78565 | 0.422285 | 2.08016 | 0.0001 | 0.01618 |
| ENSMUST00000126610 | ENSMUSG00000038900 | Rpl12 | 1242 | 141.722 | 94.2213 | 0.588938 | 0.0001 | 0.01618 |
| ENSMUST00000044355 | ENSMUSG00000041571 | Sepw1 | 736 | 417.772 | 547.636 | -0.390502 | 0.0001 | 0.01618 |
| ENSMUST00000053308 | ENSMUSG00000048424 | Ranbp3l | 3017 | 1.63158 | 2.44235 | -0.581993 | 0.0001 | 0.01618 |
| ENSMUST00000032198 | ENSMUSG00000030107 | Usp18 | 1771 | 0.459305 | 1.1285 | -1.29688 | 0.0001 | 0.01618 |
| ENSMUST00000050487 | ENSMUSG00000024610 | Cd74 | 1218 | 1.3736 | 2.74886 | -1.00087 | 0.0001 | 0.01618 |
| ENSMUST00000076657 | ENSMUSG00000035674 | Ndufa3 | 515 | 237.723 | 170.793 | 0.477038 | 0.00015 | 0.02279 |
| ENSMUST00000070112 | ENSMUSG00000026895 | Ndufa8 | 861 | 90.8273 | 69.5276 | 0.385541 | 0.00015 | 0.02279 |
| ENSMUST00000163483 | ENSMUSG00000020149 | Rab1a | 2657 | 16.0356 | 28.8646 | -0.84802 | 0.00015 | 0.02279 |
| ENSMUST00000057054 | ENSMUSG00000001493 | Meox1 | 2228 | 0.375147 | 0.824647 | -1.13632 | 0.00015 | 0.02279 |
| ENSMUST00000155120 | ENSMUSG00000057948 | Unc13d | 991 | 2.52396 | 1.1401 | 1.14652 | 0.00015 | 0.02279 |
| ENSMUST00000082432 | ENSMUSG00000007682 | Dio2 | 5813 | 17.5069 | 22.5901 | -0.367764 | 0.00015 | 0.02279 |
| ENSMUST00000089558 | ENSMUSG00000057278 | Snrpg | 1055 | 30.2101 | 21.606 | 0.483604 | 0.00015 | 0.02279 |
| ENSMUST00000044148 | ENSMUSG00000039740 | Alg2 | 3021 | 28.1938 | 36.4815 | -0.371788 | 0.00015 | 0.02279 |
| ENSMUST00000166899 | ENSMUSG00000094856 | Gm21962 | 1017 | 21.5538 | 29.1723 | -0.436654 | 0.00015 | 0.02279 |
| ENSMUST00000111336 | ENSMUSG00000058076 | Sdhc | 3171 | 13.9911 | 19.4672 | -0.476538 | 0.00015 | 0.02279 |
| ENSMUST00000102484 | ENSMUSG00000078515 | Ddi2 | 9553 | 8.30873 | 11.0824 | -0.415567 | 0.00015 | 0.02279 |
| ENSMUST00000102536 | ENSMUSG00000059291 | Rpl11 | 833 | 374.58 | 275.98 | 0.44071 | 0.00015 | 0.02279 |
| ENSMUST00000043286 | ENSMUSG00000038425 | Poli | 2591 | 2.50581 | 0.892716 | 1.489 | 0.00015 | 0.02279 |
| ENSMUST00000166898 | ENSMUSG00000091938 | Gm2564 | 707 | 66.1193 | 49.4666 | 0.418616 | 0.00015 | 0.02279 |
| ENSMUST00000115230 | ENSMUSG00000022548 | Apod | 1070 | 62.8436 | 35.8318 | 0.810527 | 0.00015 | 0.02279 |
| ENSMUST00000069180 | ENSMUSG00000055538 | Zcchc24 | 4556 | 7.67914 | 10.929 | -0.509148 | 0.00015 | 0.02279 |
| ENSMUST00000100790 | ENSMUSG00000022253 | Nadk2 | 3666 | 11.1362 | 5.72954 | 0.958766 | 0.00015 | 0.02279 |
| ENSMUST00000045376 | ENSMUSG00000039197 | Adk | 1781 | 54.8876 | 42.4967 | 0.369127 | 0.00015 | 0.02279 |
| ENSMUST00000113975 | ENSMUSG00000089774 | Slc5a3 | 10913 | 15.7962 | 12.054 | 0.390063 | 0.00015 | 0.02279 |
| ENSMUST00000063084 | ENSMUSG00000020484 | Xbp1 | 2129 | 55.7528 | 42.9004 | 0.378054 | 0.00015 | 0.02279 |
| ENSMUST00000106252 | ENSMUSG00000028654 | Mycl | 3293 | 4.70484 | 1.46103 | 1.68716 | 0.00015 | 0.02279 |

| | | | | | | | | |
|---|---|---|---|---|---|---|---|---|
| ENSMUST00000026360 | ENSMUSG00000025321 | Itgb8 | 8403 | 18.3495 | 14.4139 | 0.348272 | 0.00015 | 0.02279 |
| ENSMUST00000040912 | ENSMUSG00000036777 | Anln | 5421 | 6.31647 | 8.35374 | -0.403304 | 0.00015 | 0.02279 |
| ENSMUST00000057243 | ENSMUSG00000048572 | Tmem252 | 2462 | 0.498645 | 0.944889 | -0.922131 | 0.00015 | 0.02279 |
| ENSMUST00000021078 | ENSMUSG00000018861 | Fdxr | 1948 | 5.64346 | 3.7132 | 0.60392 | 0.00015 | 0.02279 |
| ENSMUST00000098230 | ENSMUSG00000073982 | Rhog | 1269 | 2.80934 | 4.41402 | -0.651864 | 0.00015 | 0.02279 |
| ENSMUST00000196520 | ENSMUSG00000104713 | Gbp6 | 2604 | 0.752705 | 1.30958 | -0.798951 | 0.0002 | 0.02891 |
| ENSMUST00000100171 | ENSMUSG00000026864 | Hspa5 | 2613 | 194.311 | 149.735 | 0.375959 | 0.0002 | 0.02891 |
| ENSMUST00000022416 | ENSMUSG00000021866 | Anxa11 | 2394 | 5.09607 | 3.29354 | 0.629745 | 0.0002 | 0.02891 |
| ENSMUST00000087714 | ENSMUSG00000067455 | Hist1h4j | 774 | 27.6825 | 20.0283 | 0.466933 | 0.0002 | 0.02891 |
| ENSMUST00000135941 | ENSMUSG00000006412 | Pfdn2 | 1735 | 13.954 | 19.6005 | -0.490217 | 0.0002 | 0.02891 |
| ENSMUST00000108935 | ENSMUSG00000074521 | Gm14327 | 1451 | 7.88642 | 5.66907 | 0.476257 | 0.0002 | 0.02891 |
| ENSMUST00000028162 | ENSMUSG00000026820 | Ptges2 | 4978 | 7.64738 | 5.47653 | 0.481702 | 0.0002 | 0.02891 |
| ENSMUST00000022622 | ENSMUSG00000059456 | Ptk2b | 4034 | 64.6566 | 44.8887 | 0.526446 | 0.0002 | 0.02891 |
| ENSMUST00000154650 | ENSMUSG00000013523 | Bcas1 | 2222 | 14.8288 | 20.0485 | -0.435097 | 0.0002 | 0.02891 |
| ENSMUST00000170715 | ENSMUSG00000044533 | Rps2 | 998 | 23.6517 | 36.76 | -0.63619 | 0.0002 | 0.02891 |
| ENSMUST00000039784 | ENSMUSG00000036138 | Acaa1a | 1762 | 16.6611 | 8.00985 | 1.05664 | 0.0002 | 0.02891 |
| ENSMUST00000080322 | ENSMUSG00000052776 | Oas1a | 1885 | 0.334109 | 0.841437 | -1.33254 | 0.0002 | 0.02891 |
| ENSMUST00000127748 | ENSMUSG00000043496 | Tril | 5363 | 5.04737 | 3.76193 | 0.424059 | 0.0002 | 0.02891 |
| ENSMUST00000035113 | ENSMUSG00000032526 | Deb1 | 517 | 80.2409 | 59.9002 | 0.421777 | 0.0002 | 0.02891 |
| ENSMUST00000102840 | ENSMUSG00000076441 | Ass1 | 1696 | 17.544 | 23.1705 | -0.401316 | 0.0002 | 0.02891 |
| ENSMUST00000071135 | ENSMUSG00000062591 | Tubb4a | 2228 | 86.6397 | 110.242 | -0.347568 | 0.0002 | 0.02891 |
| ENSMUST00000109279 | ENSMUSG00000018459 | Slc13a3 | 3131 | 1.61907 | 0.513147 | 1.65773 | 0.0002 | 0.02891 |
| ENSMUST00000136312 | ENSMUSG00000008348 | Ubc | 2618 | 92.0182 | 115.448 | -0.327247 | 0.0002 | 0.02891 |
| ENSMUST00000016640 | ENSMUSG00000016496 | Cd274 | 3622 | 0.589914 | 0.95587 | -0.696309 | 0.0002 | 0.02891 |
| ENSMUST00000072079 | ENSMUSG00000061024 | Rrs1 | 2048 | 6.58605 | 9.12347 | -0.470169 | 0.0002 | 0.02891 |
| ENSMUST00000007139 | ENSMUSG00000006941 | Eif1b | 917 | 100.829 | 77.5519 | 0.378671 | 0.00025 | 0.03454 |
| ENSMUST00000089752 | ENSMUSG00000053754 | Chd8 | 8190 | 3.70081 | 6.71823 | -0.860241 | 0.00025 | 0.03454 |
| ENSMUST00000029451 | ENSMUSG00000027858 | Tspan2 | 4484 | 13.4033 | 17.3483 | -0.372207 | 0.00025 | 0.03454 |
| ENSMUST00000029271 | ENSMUSG00000027716 | Trpc3 | 3694 | 4.31024 | 5.78166 | -0.423717 | 0.00025 | 0.03454 |
| ENSMUST00000021203 | ENSMUSG00000020843 | Timm22 | 2817 | 14.4595 | 10.9161 | 0.405557 | 0.00025 | 0.03454 |
| ENSMUST00000166791 | ENSMUSG00000042502 | Cd2bp2 | 3284 | 12.5864 | 7.63879 | 0.720448 | 0.00025 | 0.03454 |
| ENSMUST00000145910 | ENSMUSG00000024077 | Strn | 8629 | 20.1398 | 15.68 | 0.361124 | 0.00025 | 0.03454 |
| ENSMUST00000055389 | ENSMUSG00000047434 | Xxylt1 | 2987 | 4.11637 | 5.67191 | -0.462462 | 0.00025 | 0.03454 |
| ENSMUST00000053764 | ENSMUSG00000044167 | Foxo1 | 8665 | 1.77147 | 2.36563 | -0.417276 | 0.00025 | 0.03454 |
| ENSMUST00000001825 | ENSMUSG00000001774 | Chordc1 | 2205 | 46.5842 | 36.4915 | 0.35228 | 0.00025 | 0.03454 |
| ENSMUST00000096259 | ENSMUSG00000071658 | Gng3 | 1699 | 31.7204 | 44.8122 | -0.498483 | 0.00025 | 0.03454 |
| ENSMUST00000067924 | ENSMUSG00000054720 | Lrrc8c | 6976 | 3.33745 | 4.42946 | -0.408384 | 0.00025 | 0.03454 |
| ENSMUST00000163582 | ENSMUSG00000059895 | Ptp4a3 | 3808 | 3.97246 | 0.681066 | 2.54417 | 0.00025 | 0.03454 |
| ENSMUST00000176010 | ENSMUSG00000093674 | Rpl41 | 582 | 984.151 | 615.812 | 0.676389 | 0.00025 | 0.03454 |
| ENSMUST00000102844 | ENSMUSG00000020460 | Rps27a | 637 | 139.559 | 80.7438 | 0.789455 | 0.00025 | 0.03454 |
| ENSMUST00000032944 | ENSMUSG00000030703 | Gdpd3 | 1148 | 5.25516 | 3.10452 | 0.759366 | 0.00025 | 0.03454 |
| ENSMUST00000040077 | ENSMUSG00000033020 | Polr2f | 543 | 64.0702 | 47.605 | 0.428541 | 0.00025 | 0.03454 |
| ENSMUST00000106355 | ENSMUSG00000045268 | Zfp691 | 1588 | 1.35804 | 0.622856 | 1.12455 | 0.00025 | 0.03454 |
| ENSMUST00000061169 | ENSMUSG00000047658 | Gal3st3 | 2421 | 6.16062 | 8.33777 | -0.436585 | 0.00025 | 0.03454 |
| ENSMUST00000035625 | ENSMUSG00000039519 | Cyp7b1 | 2166 | 10.5074 | 7.62523 | 0.462552 | 0.00025 | 0.03454 |
| ENSMUST00000002379 | ENSMUSG00000002308 | Cd320 | 2209 | 4.33375 | 2.31034 | 0.907512 | 0.00025 | 0.03454 |
| ENSMUST00000008966 | ENSMUSG00000008822 | Acyp1 | 648 | 47.6244 | 35.5216 | 0.423004 | 0.0003 | 0.03991 |
| ENSMUST00000046644 | ENSMUSG00000042043 | Tbca | 503 | 212.397 | 166.256 | 0.353359 | 0.0003 | 0.03991 |
| ENSMUST00000187137 | ENSMUSG00000036634 | Mag | 2434 | 22.191 | 30.1232 | -0.440902 | 0.0003 | 0.03991 |
| ENSMUST00000048781 | ENSMUSG00000040618 | Pck2 | 3400 | 6.94631 | 5.25421 | 0.402775 | 0.0003 | 0.03991 |
| ENSMUST00000077741 | ENSMUSG00000060681 | Slc9a6 | 4976 | 18.1373 | 11.9578 | 0.601013 | 0.0003 | 0.03991 |
| ENSMUST00000056153 | ENSMUSG00000044788 | Fads6 | 4951 | 4.80383 | 6.31882 | -0.395468 | 0.0003 | 0.03991 |
| ENSMUST00000022145 | ENSMUSG00000021643 | Serf1 | 601 | 24.6667 | 15.0113 | 0.716516 | 0.0003 | 0.03991 |
| ENSMUST00000101800 | ENSMUSG00000075705 | Msrb1 | 1211 | 1.4731 | 3.5774 | -1.28006 | 0.0003 | 0.03991 |
| ENSMUST00000005620 | ENSMUSG00000005483 | Dnajb1 | 2263 | 40.8973 | 31.9944 | 0.354186 | 0.0003 | 0.03991 |
| ENSMUST00000027467 | ENSMUSG00000026249 | Serpine2 | 2210 | 65.5944 | 50.4842 | 0.377741 | 0.0003 | 0.03991 |
| ENSMUST00000069451 | ENSMUSG00000055737 | Ghr | 4175 | 2.49909 | 1.70992 | 0.547474 | 0.0003 | 0.03991 |
| ENSMUST00000108980 | ENSMUSG00000078878 | Gm14305 | 1167 | 23.4126 | 15.9274 | 0.555774 | 0.0003 | 0.03991 |
| ENSMUST00000024802 | ENSMUSG00000024007 | Ppil1 | 1434 | 11.326 | 8.26976 | 0.453725 | 0.0003 | 0.03991 |
| ENSMUST00000146824 | ENSMUSG00000031683 | Lsm6 | 541 | 37.2603 | 16.3913 | 1.18471 | 0.0003 | 0.03991 |
| ENSMUST00000183338 | ENSMUSG00000037174 | Elf2 | 1083 | 43.8741 | 29.0886 | 0.592916 | 0.0003 | 0.03991 |
| ENSMUST00000155802 | ENSMUSG00000038147 | Cd84 | 3291 | 0.239949 | 0.601229 | -1.32519 | 0.0003 | 0.03991 |
| ENSMUST00000060214 | ENSMUSG00000050854 | Tmem125 | 1864 | 1.71869 | 2.66569 | -0.633194 | 0.0003 | 0.03991 |
| ENSMUST00000205741 | ENSMUSG00000013465 | Nelfb | 2636 | 17.1853 | 12.69 | 0.437483 | 0.0003 | 0.03991 |
| ENSMUST00000067048 | ENSMUSG00000022262 | Dnah5 | 15630 | 0.979096 | 1.31997 | -0.430986 | 0.0003 | 0.03991 |
| ENSMUST00000030626 | ENSMUSG00000028822 | Tmem50a | 1204 | 59.5914 | 76.0637 | -0.352103 | 0.0003 | 0.03991 |
| ENSMUST00000078676 | ENSMUSG00000063882 | Uqcrh | 547 | 572.299 | 443.527 | 0.367749 | 0.0003 | 0.03991 |
| ENSMUST00000170953 | ENSMUSG00000090862 | Rps13 | 589 | 267.831 | 208.331 | 0.362445 | 0.0003 | 0.03991 |
| ENSMUST00000029402 | ENSMUSG00000027822 | Slc33a1 | 3150 | 13.2838 | 8.93294 | 0.572458 | 0.0003 | 0.03991 |
| ENSMUST00000043148 | ENSMUSG00000036402 | Gng12 | 4248 | 11.2786 | 8.5516 | 0.399319 | 0.00035 | 0.04521 |
| ENSMUST00000034728 | ENSMUSG00000032198 | Dock6 | 6556 | 0.617658 | 0.920785 | -0.576057 | 0.00035 | 0.04521 |
| ENSMUST00000033342 | ENSMUSG00000031029 | Eif3f | 2514 | 26.1737 | 20.2374 | 0.371096 | 0.00035 | 0.04521 |
| ENSMUST00000125077 | ENSMUSG00000052407 | Ccdc171 | 4231 | 1.29963 | 0.787197 | 0.723309 | 0.00035 | 0.04521 |
| ENSMUST00000039758 | ENSMUSG00000035885 | Cox8a | 545 | 1352.61 | 1072.99 | 0.334115 | 0.00035 | 0.04521 |
| ENSMUST00000067925 | ENSMUSG00000054717 | Hmgb2 | 2692 | 1.00867 | 1.56782 | -0.636308 | 0.00035 | 0.04521 |
| ENSMUST00000177921 | ENSMUSG00000096808 | AC132444.6 | 1500 | 1.46109 | 2.5448 | -0.800512 | 0.00035 | 0.04521 |
| ENSMUST00000129829 | ENSMUSG00000039997 | Ifi203 | 3054 | 0.118924 | 0.416118 | -1.80695 | 0.00035 | 0.04521 |
| ENSMUST00000043675 | ENSMUSG00000038717 | Atp5l | 511 | 612.66 | 484.633 | 0.338194 | 0.00035 | 0.04521 |
| ENSMUST00000169927 | ENSMUSG00000042429 | Adora1 | 4581 | 5.35874 | 9.97924 | -0.897037 | 0.00035 | 0.04521 |
| ENSMUST00000026565 | ENSMUSG00000025492 | Ifitm3 | 678 | 3.92549 | 6.53203 | -0.734659 | 0.00035 | 0.04521 |

| | | | | | | | | |
|---|---|---|---|---|---|---|---|---|
| ENSMUST00000112359 | ENSMUSG00000042328 | Hps4 | 3345 | 3.33248 | 1.75422 | 0.925769 | 0.00035 | 0.04521 |
| ENSMUST00000038194 | ENSMUSG00000022360 | Atad2 | 5683 | 3.80782 | 2.84409 | 0.420995 | 0.00035 | 0.04521 |
| ENSMUST00000020970 | ENSMUSG00000020641 | Rsad2 | 3775 | 0.39366 | 0.681637 | -0.792054 | 0.00035 | 0.04521 |
| ENSMUST00000008830 | ENSMUSG00000094441 | Zfp955a | 1638 | 18.3613 | 14.0446 | 0.386649 | 0.00035 | 0.04521 |
